# piRNA processing by a trimeric Schlafen-domain nuclease

**DOI:** 10.1101/2023.01.19.524756

**Authors:** Nadezda Podvalnaya, Alfred W. Bronkhorst, Raffael Lichtenberger, Svenja Hellmann, Emily Nischwitz, Torben Falk, Emil Karaulanov, Falk Butter, Sebastian Falk, René F. Ketting

## Abstract

Transposable elements are genomic parasites that expand within and spread between genomes^1^. Piwi proteins control transposon activity, notably in the germline^2,3^. These proteins recognize their targets through small RNA co-factors named piRNAs, making piRNA biogenesis a key specificity-determining step in this crucial genome immunity system. While the processing of piRNA precursors is an essential step in this process, many molecular details of this process remain unknown. We identify a novel endoribonuclease, PUCH, that initiates piRNA processing in the nematode *Caenorhabditis elegans*. Genetic and biochemical studies show that PUCH, a trimer of Schlafen-like-domain proteins (SLFL proteins), executes 5’-end piRNA precursor cleavage. PUCH-mediated processing strictly requires an m7G-Cap and a uracil at position three. We also demonstrate how PUCH interacts with PETISCO, a complex that binds piRNA precursors^4^, and that this interaction enhances piRNA production *in vivo*. The identification of PUCH completes the repertoire of *C. elegans* piRNA biogenesis factors and uncovers a novel type of RNA endonuclease formed by three SLFL proteins. Mammalian Schlafen (Slfn) genes have been associated with immunity responses^5^, exposing a thus far unknown molecular link between immune responses in mammals and deeply conserved RNA-based mechanisms that control transposable elements.

## Introduction

Transposable elements (TEs) are segments of DNA that can independently multiply and move within, and sometimes between genomes^1^. Being found in virtually all genomes analyzed to date, transposons are clearly highly successful, and hence their control, especially in the germ cells, is an essential process. Interestingly, TEs can mutate to avoid defense systems, and in turn, defense systems can adapt to such changes, resulting in a molecular arms race, leading to rapid diversification between species^6^. Small RNA-driven gene regulatory pathways represent one of the mechanisms through which transposable elements are controlled^2,3^. In the germ cells of animals, Argonaute proteins of the Piwi clade interact with so-called piRNAs to control transposons, but also host genes^7^. This regulatory process is essential for germ cell function and fertility. Consistent with a role in TE defense, piRNA pathways are characterized by many species-specific factors, even though piRNA pathways also share deeply conserved concepts^2,3^.

The production of piRNAs is essential, as the piRNAs portfolio defines the target range and specificity of the Piwi-piRNA pathway. Mature piRNAs are generated from longer, single-stranded piRNA precursor molecules^2,3,8^. In all systems analyzed, this process is started by a nucleolytic cleavage, which defines the 5’-end of a new piRNA. In *Drosophila* and mouse, Piwi mediated cleavage (slicer activity) can generate such a 5’-end, which then is bound by a new Piwi protein. The net effect of this process is the amplification of piRNAs through Piwi cleavage. Moreover, the Piwi protein that binds to the newly generated 5’-end recruits an endonuclease named Zucchini (Zuc), or PhospholipaseD6 (PLD6)^9–12^. Zuc induces an additional break, and thus an additional 5’-end, downstream of the bound Piwi protein, which can again be bound by a Piwi protein ^13,14^. This process not only amplifies but also diversifies piRNA populations. The current model suggests that after 5’-end processing the 3’-end is formed following binding to a Piwi protein. This step involves trimming by 3’-5’ exoribonuclease activity and methylation of the 2’ OH at the 3’-end. In *C. elegans* this is done by PARN-1^15^ and HENN-1 respectively^16–18^.

Interestingly, not all animals rely on Zuc/PLD6 and/or Piwi for piRNA biogenesis. Notably, *C. elegans* lacks a Zuc homolog. Furthermore, the slicer activity of the *C. elegans* Piwi homolog (PRG-1^19–21^) is not needed for piRNA production^22^, making it a great unknown how piRNA 5’-ends are generated. In this nematode, piRNA precursors are transcribed from short genes, each encoding one piRNA, which in *C. elegans* is also named 21U RNA^23^. The precursors are roughly 27-29 nucleotides long and carry a 5’-Cap (**Fig. 1a**)^24^. Since these precursors are not trans-spliced, this Cap is most likely an m7-G Cap. We note that in contrast to many other animals, including mammals, *C. elegans* does not place m7-G Caps on canonical mRNAs, but 2,2,7-trimethyl-G Caps (TMG-Cap) through a process of 5’-end trans-splicing^25^. It is thus possible that the m7-G Cap can be a feature that helps distinguish between piRNA precursors and mRNAs. Following transcription by specialized machinery^26,27^, the precursors are bound by PETISCO, a cytoplasmic protein complex consisting of PID-3, ERH-2, TOFU-6 and IFE-3^4,28–30^. While PETISCO has been implicated in precursor stabilization and is required for piRNA production, it contains no nucleases. Hence, the nuclease that mediates 5’ precursor processing and how it interact with PETISCO remains unidentified.

**Fig. 1.**
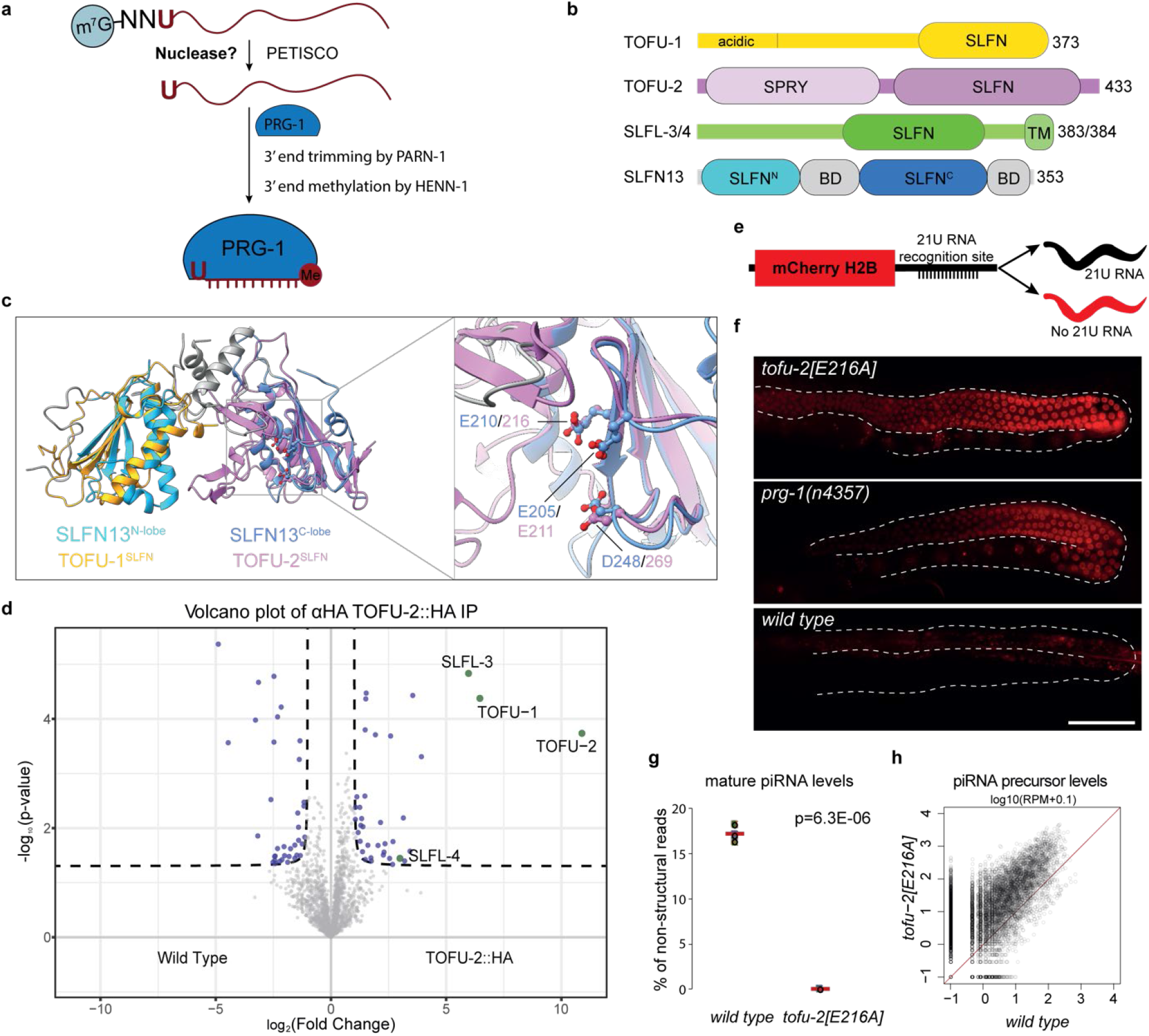
Identification of the catalytic center of TOFU-2. **a**, Model of piRNA (21U RNA) formation in *C. elegans*. Individually transcribed piRNA precursors of ∼27-29 nucleotides in length are stabilized by the PETISCO complex. Following removal of the 5’ Cap and two nucleotides, the intermediates are loaded onto PRG-1, followed by trimming and methylation of the 3’-end by PARN-1 and HENN-1, respectively. The nuclease that is responsible for 5’-end maturation of piRNAs has remained unknown thus far. **b**, Schematic domain organization of TOFU-1, TOFU-2 and SLFL-3/4. Lines indicate low-complexity regions and rounded rectangles indicate predicted folded domains. BD: bridging domain, TM: transmembrane domain. **c**, Superposition of the TOFU-1 and TOFU-2 SLFN domains onto the crystal structure of the N-terminal SLFN13 endoribonuclease domain. The domains are colored as in panel (b). The SLFN domains of TOFU-1 is shown in yellow, of TOFU-2 in purple and of SLFN13 in light and dark blue. The SLFN13 bridging domains in grey. The zoom-in view shows the active site of SLFN13 and involved residues are shown as sticks. **d**, Volcano plot representing label-free proteomic quantification of TOFU-2::HA (n=4) and wild type (n=4, WT) immunoprecipitations from young adult extracts. The X-axis represents the median fold enrichment of individual proteins in wild type (WT) versus the TOFU-2::HA mutant strain. The Y-axis indicates −log10(P-value) calculated using Welch t-test. Dashed lines represent enrichment thresholds at p-value = 0.05 and fold change > 2, c = 0.05. Each dot represents an enriched (blue/green) or quantified (grey) proteins. **e**, Schematic representation of the mCherry::H2B piRNA sensor. In presence of piRNAs, mCherry will be repressed, whereas in the absence of piRNA mCherry reporter will be expressed resulting in red fluorescent nuclei in the germline. **f**, Widefield fluorescent microscopy of adult hermaphrodites carrying the piRNA sensor in three genetic backgrounds: *tofu-2[E216A]* on top, *prg-1(n4357)* in the middle, wild type at the bottom. The germlines are outlined by a white dashed line. Scale bar – 50 µm. **g**, Total piRNA levels in wild type and *tofu-2[E216A]*-mutant young adult hermaphrodites (n = 3). Red lines depict group means and P-values were calculated using two-tailed unpaired t-test. **h**, Scatter plot depicting the relative abundance of piRNA precursors from individual loci in *tofu-2[E216A]*-mutant versus wild type young adult hermaphrodites. RPM: Reads per million non-structural sRNA reads.

### TOFU-2 interacts with TOFU-1 and is a potential nuclease

A genome-wide RNAi screen identified the proteins TOFU-1 and TOFU-2 as factors necessary for piRNA accumulation^31^. Loss of these factors also triggered piRNA precursor accumulation, suggesting they may play a role in piRNA 5’-end processing. However, domain annotations at that time did not reveal potential nuclease domains. Using structure-based homology searches (HHPRED)^32^ and AlphaFold2 (AF2)^33^, we detected homology between TOFU-1 and TOFU-2, and the rat ribonuclease Schlafen13 (SLFN13)^34^, but also human Schlafen5^35^ and Schlafen12^36^. This identified the presence of a potential SLFN-fold in both TOFU-1 and TOFU-2 (**Fig. 1b,c**). Interestingly, whereas in mammalian Schlafen proteins two SLFN-folds come together to form the nuclease domain^34–36^, in TOFU-1 and TOFU-2 only one SLFN-fold could be identified. Hence, we hypothesized that TOFU-1 and TOFU-2 may interact to form a functional nuclease. To test this hypothesis, we tagged endogenous TOFU-2 with a Human influenza hemagglutinin (HA)-tag and used immuno-precipitation (IP) followed by quantitative mass spectrometry (MS) to identify TOFU-2 interacting proteins. Indeed, TOFU-1 was found to interact with TOFU-2 (**Fig.1d; Table S1**). Potential catalytic residues were identified within TOFU-2, but not TOFU-1 (**Fig.1c, Suppl. Fig.1a**). Thus, we engineered a *C. elegans tofu-2* mutant in which we changed one of the potential catalytic residues (glutamic acid 216) to alanine (E216A). This mutation neither affects TOFU-2 abundance nor interaction with TOFU-1, as judged by Western blot analysis and IP-MS analysis (**Suppl. Fig.1c,d; Table S2**). Next, we tested piRNA silencing activity in this mutant, using a so-called piRNA-sensor (a germline-expressed transgene that is silenced through piRNA activity^22^) (**Fig.1e**). This revealed that *tofu-2(e216a)* mutants de-silence the piRNA-sensor to a similar extent as *prg-1* mutants (**Fig.1f**). Sequencing of piRNAs and piRNA precursors showed that *tofu-2(e216a)* mutants lost almost all mature piRNAs, while accumulating precursors (**Fig.1g, h, Suppl. Fig.1e**), suggesting that a TOFU-1:TOFU-2 complex could be the nuclease that processes piRNA precursors.

### TOFU-1 and TOFU-2 interact with SLFL-3 or SLFL-4 to form a trimeric complex

Next, we heterologously expressed TOFU-1 and TOFU-2 in BmN4 cells, a silk-moth derived cell culture system that expressed these proteins well, and found that TOFU-1 and TOFU-2 co-immunoprecipitate (coIP) each other (**Fig.2a, lane 4**). However, incubating the coIPs with a synthetic piRNA precursor did not result in precursor cleavage (see next section), suggesting that our experimental conditions might lack an essential cofactor.

**Fig. 2.**
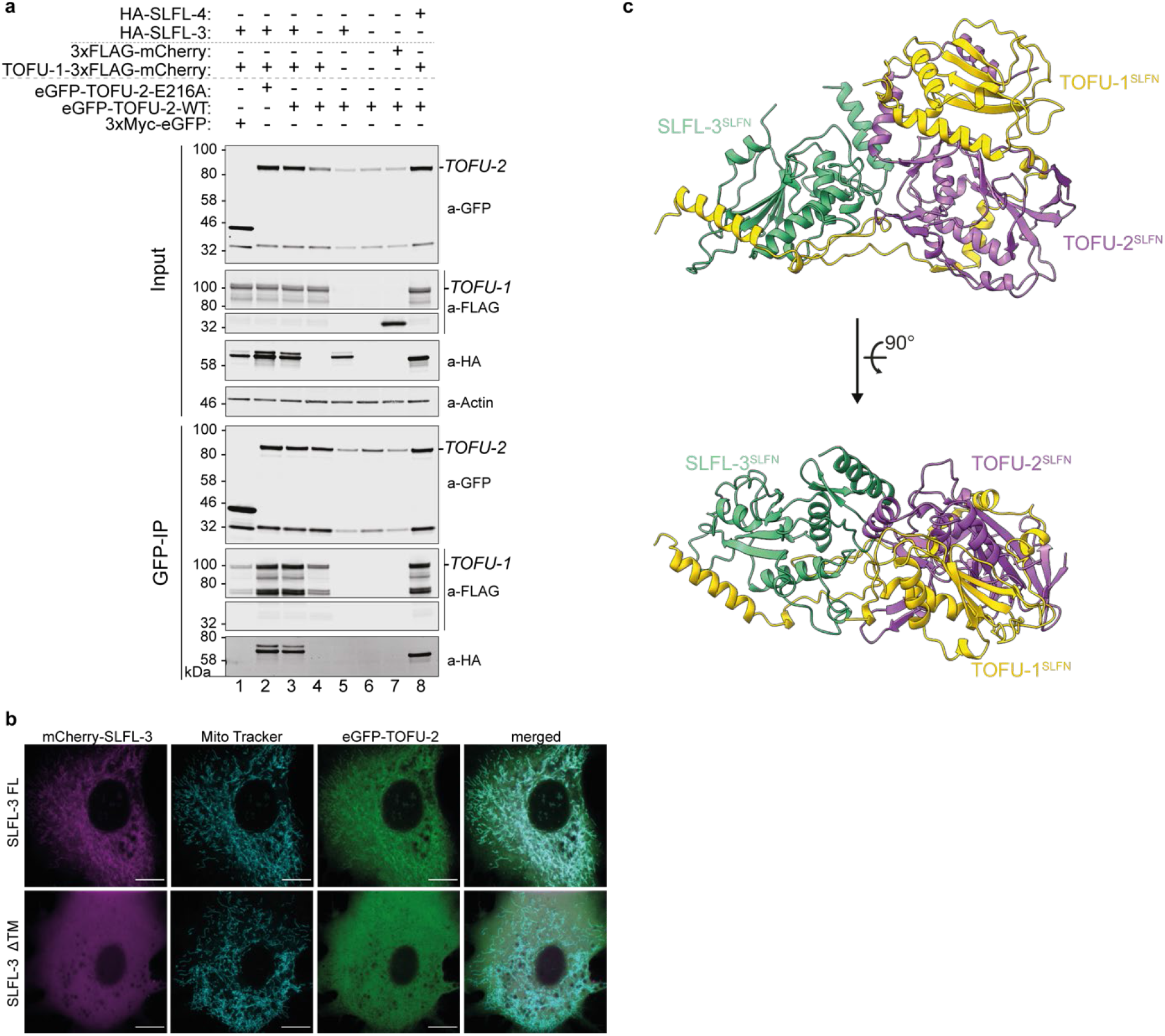
TOFU-1, TOFU-2 and SLFL-3/4 form a mitochondria-bound complex. **a**, Anti-GFP immunoprecipitation from BmN4 cell lysates made from cells that were transfected with the indicated constructs. eGFP-TOFU-2 was immunoprecipitated, followed by Western blot detection of TOFU-1 (FLAG), SLFL-3 (HA) or SLFL-4 (HA). Expression of 3xMyc-eGFP and of 3xFLAG-mCherry served as negative controls. **b**, Single-plane confocal micrographs of BmN4 cells transfected with eGFP-TOFU-2 and full-length mCherry-SLFL-3 (top) or mCherry-SLFL-3 ΔTM (bottom). TOFU-1 was also transfected but was not tagged with a fluorescent protein. Mitochondria were stained with Mito Tracker. Scale bars – 10 µm. **c**, AlphaFold predicted structure of a minimal trimeric TOFU-1, TOFU-2, SLFL-3 complex. The best of five predicted models is shown as cartoon in two different orientations. TOFU-1 is shown in yellow, TOFU-2 in purple and SLFL-3 in green.

The TOFU-2 IP-MS experiments, in addition to TOFU-1, also identified the proteins C35E7.8 and F36H12.2 (**Fig.1d, Suppl. Fig.1a,b**). These two proteins are 90% identical at the amino-acid level (**Suppl. Fig.2a**) and hence may function redundantly. Analysis with AF2^33,37^ revealed that these two proteins also contain a single potential SLFN-like fold (**Suppl. Fig.2b**). Therefore, we propose the name “SLFN-like”, or SLFL, for this group of proteins that only contain a single SLFN-fold, with TOFU-1, TOFU-2, C35E7.8 and F36H12.2 corresponding to SLFL-1, SLFL-2, SLFL-3 and SLFL-4 respectively. Interestingly, SLFL-3 was identified in the same study that identified TOFU-1 and TOFU-2, but its RNAi-mediated knock-down triggered a relatively weak reduction in piRNA levels and was not investigated further^31^. We generated a *slfl-3* deletion mutant (**Suppl. Fig.2c**) and found that this allele triggers reduced piRNA activity, as evidenced by a mild activation of the piRNA sensor (**Suppl. Fig.2d**). In addition to the SLFN-like fold, SLFL-3 and SLFL-4 also contain a predicted transmembrane (TM) helix (**Suppl. Fig.2a,e**), a feature that is also present in mammalian and *Drosophila* Zuc^38,39^. By transfecting BmN4 cells with TOFU-1, eGFP-TOFU-2 and mCherry-SLFL-3 carrying or lacking the TM, we could show that the TM of SLFL-3 localizes to mitochondria (**Fig.2b**). Interestingly, TOFU-2 colocalizes with SLFL-3 suggesting they form a complex.

We used AF2^33,40^ to predict how these four SLFL proteins may interact with each other. This revealed that a trimeric combination of TOFU-1, TOFU-2 and either SLFL-3 or SLFL-4 yielded the best predictions, in which the three SLFN domains were found to interact with each other (**Fig. 2c, Suppl. Fig.3**). Further fine-tuning of the procedure produced a high-confidence model of TOFU-1, TOFU-2 and SLFL-3 (**Fig. 2c, Suppl. Fig.4**) suggesting that the active nuclease may be a trimeric complex. This prompted us to co-express TOFU-1, TOFU-2 and either SLFL-3 or SLFL-4 in BmN4 cells and to test their interaction through coIP experiments. Indeed, these experiments support the idea of a trimer. For instance, TOFU-2 is expressed at higher levels if co-expressed with TOFU-1 and SLFL-3 and TOFU-2 only coIPs with SLFL-3 in the presence of TOFU-1 (**Fig.2a**).

**Fig. 3.**
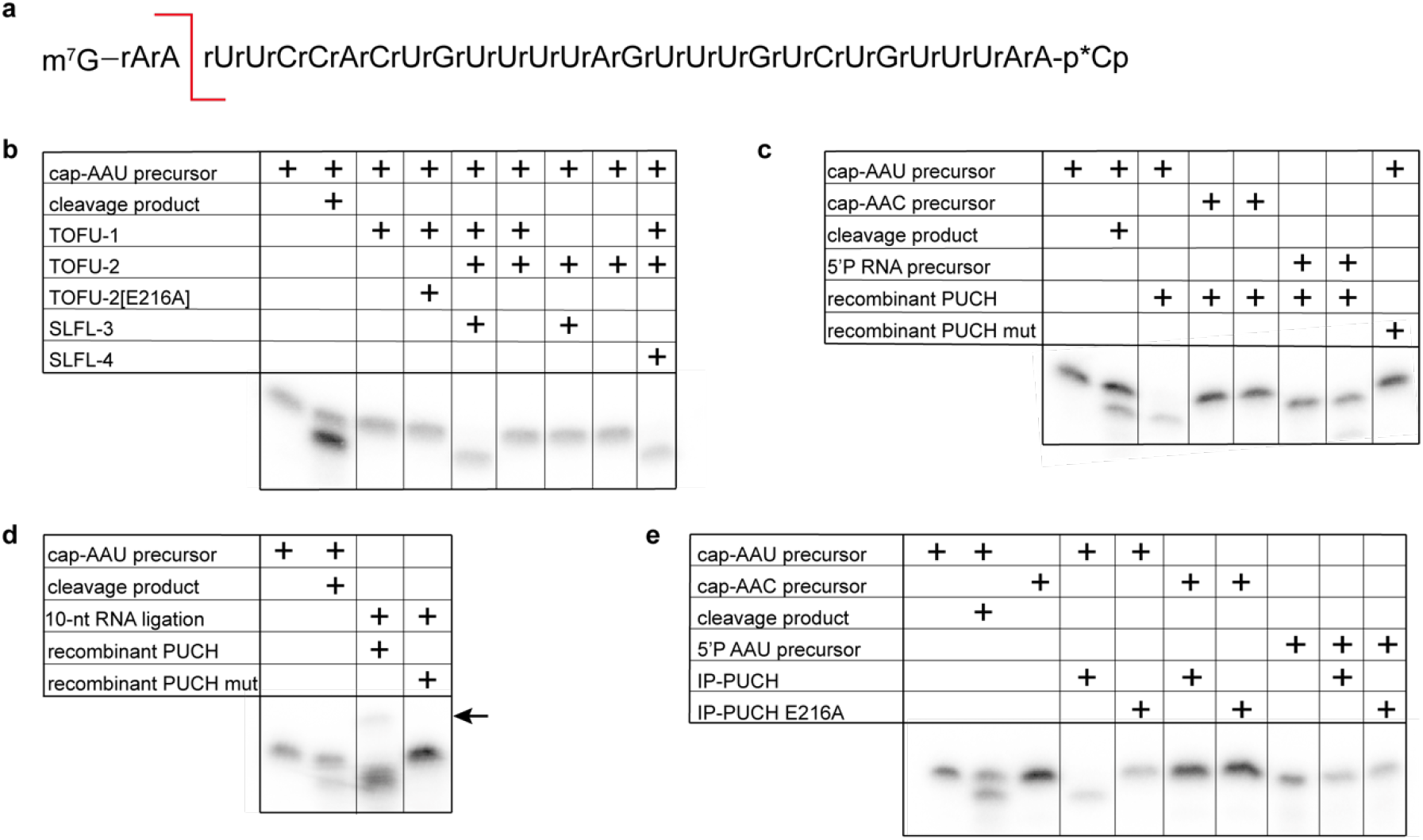
PUCH is a Cap- and sequence-specific endoribonuclease. **a**, Sequence of the synthetic piRNA precursor used in the assay. Red line indicates the expected cleavage position. Both precursor and expected cleavage product were run in the two left-most lanes of every gel to mark where these molecules are to be expected. **b**, *In vitro* cleavage assay of the piRNA precursor using GFP-IP material from BmN4 cell extracts. Cells were transfected with eGFP-TOFU-2, TOFU-1, SLFL-3 or SLFL-4 in various combinations, as indicated. **c**, Cleavage assays with recombinant PUCH and different RNA substrates. ‘Recombinant PUCH Mut’ reflects a complex in which TOFU-2 was mutated in the catalytic site (TOFU-2[E216A]). **d**, RNA obtained from a cleavage reaction (using either wild-type or TOFU-2[E216A] mutant PUCH complex) was ligated to a 10 nucleotide long 5’OH-containing RNA adapter. The ligation product is indicated with an arrow. **e**, *In vitro* cleavage assay on different types of RNA substrate using the PUCH complex retrieved from BmN4 cells by immunoprecipitation.

**Fig. 4.**
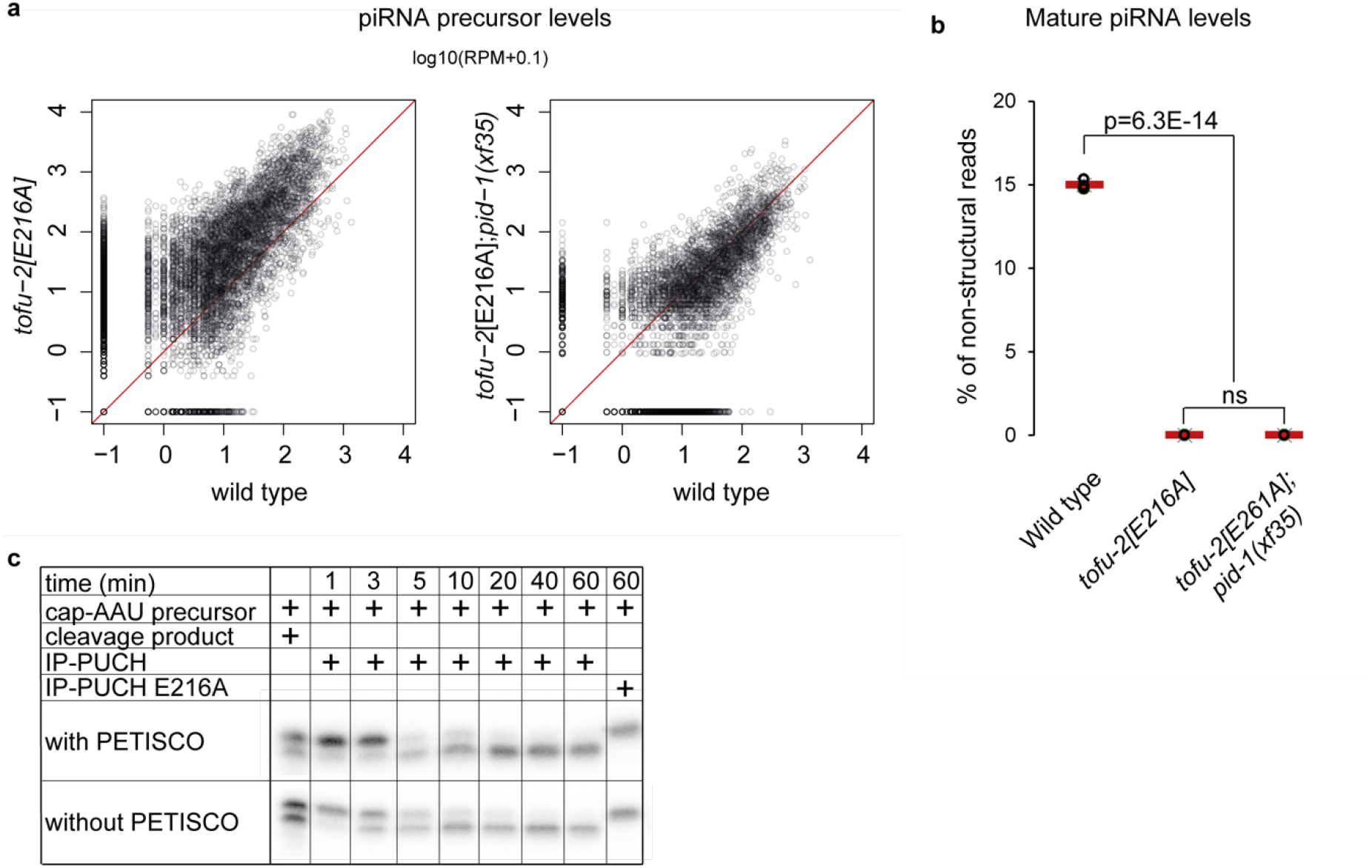
PETISCO is necessary for piRNA precursor accumulation *in vivo* and does not interfere with PUCH-mediated precursor cleavage. **a**, Scatter plots depicting the relative abundance of individual piRNA precursors in *tofu-2[E216A]*-mutant (left) and *tofu-2[E216A];pid-1(xf35)-*double mutant (right) versus wild type young adult hermaphrodites. RPM: Reads per million non-structural sRNA reads. **b**, Total piRNA levels in wild type, *tofu-2[E216A]*-mutant and *tofu-2[E216A];pid-1(xf35)-* double mutant young adult hermaphrodites (n=3). Red lines depict group means and P-values were calculated using ANOVA test followed by Tukey HSD. **c**, *In vitro* piRNA precursor cleavage assay either in presence or absence of the PETISCO complex in a time-series. PUCH in this experiment was isolated from BmN4 cell extracts using immune-precipitations.

We also assessed the interactions between TOFU-1, TOFU-2 and SLFL-3 through heterologous expression and coIP experiments in *E. coli* (**Suppl. Fig.5a**). While the TOFU-2 SPRY domain did not display strong interactions (**Suppl. Fig.5b**), TOFU-1 and SLFL-3 interacted with their SLFN-domain directly with the TOFU-2 SLFN domain (**Suppl. Fig.5c-d**), and a complex containing all three proteins could be readily identified (**Suppl. Fig.5e**). These findings are in line with the AF2 model (**Suppl. Fig.4**) and the coIPs from BmN4 cells (**Fig.2a**).

**Fig. 5.**
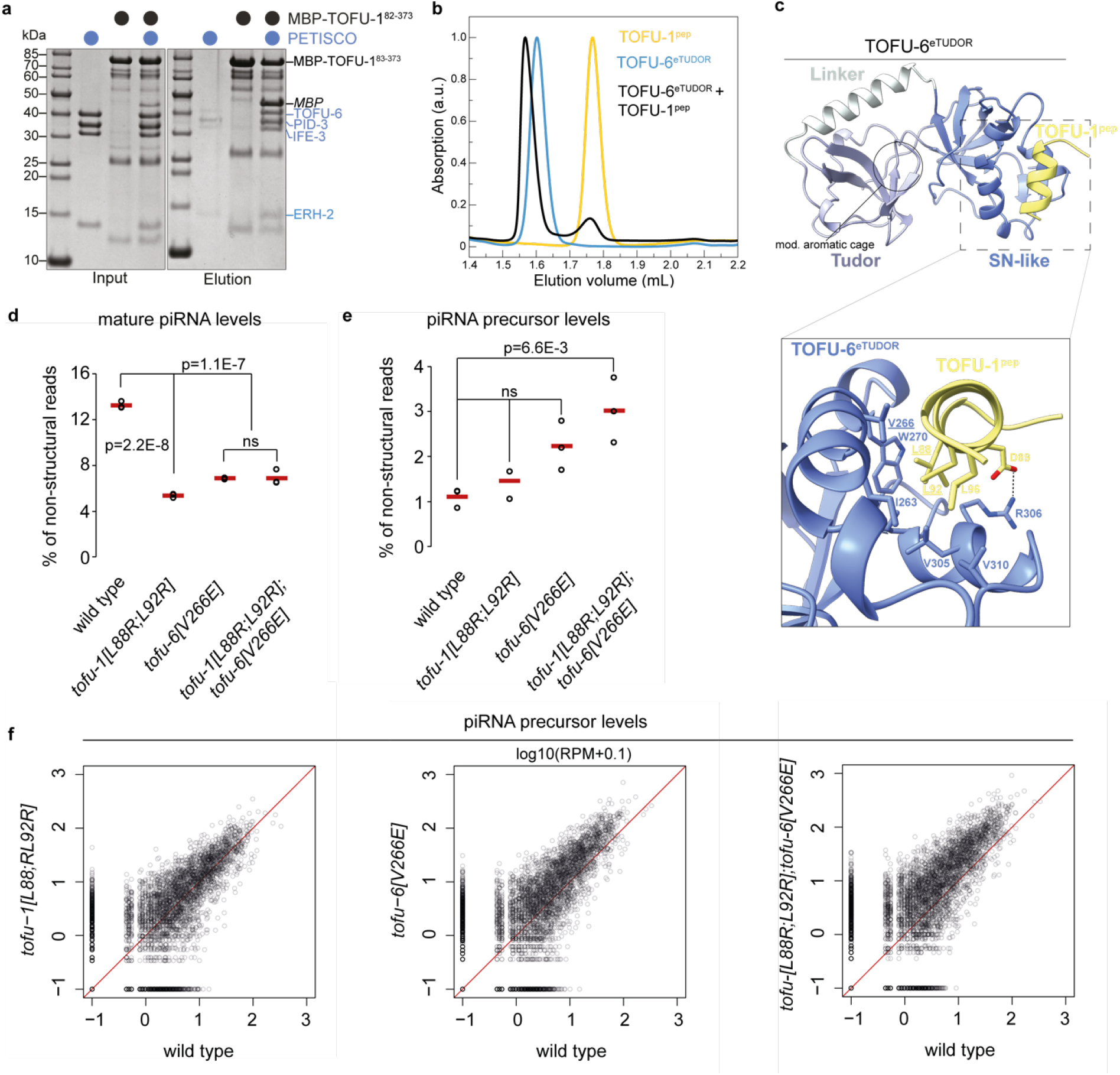
TOFU-6 from PETISCO interacts with PUCH via TOFU-1. **a**, Analysis of the interaction between TOFU-1 and PETISCO by amylose pull-down assays. Purified MBP-tagged TOFU-1^82-373^ was incubated with excess PETISCO. Input and elution fractions were analyzed by SDS-PAGE followed by Coomassie staining. **b**, Purified TOFU-6^eTUDOR^, TOFU-1^82-113^ (=TOFU-1^pep^) and a mixture thereof were subjected to size exclusion chromatography. Chromatograms: TOFU-6^eTUDOR^ (blue), TOFU-1^pep^ (yellow) and TOFU-6^eTUDOR^ + TOFU-1^pep^ (black). **c**, Crystal structure of the TOFU-6^eTUDOR^–TOFU-1^pep^ complex shown as cartoon. The TOFU-6^eTUDOR^ domain is shown in different shades of blue and TOFU-1^pep^ in yellow. The zoom-in view shows the interaction interface; involved residues are shown as sticks. **d and e**, Total mature (**d**) and precursor (**e**) piRNA levels in wild-type, *tofu-1[L88R;L92R], tofu-6[V266E]* and *tofu-1[L88R;L92R];tofu-6[V266E]*-double mutant young adult hermaphrodites (n = 3). Red lines depict group means and P-values were calculated using ANOVA test followed by Tukey HSD. **f**, Scatter plots depicting the relative abundance of precursors from individual piRNA loci in *tofu-1[L88R;L92R]* (left), *tofu-6[V266E]* (middle) and *tofu-1[L88R;L92R];tofu-6[V266E]*-double mutant (right) versus wild-type young adult hermaphrodites. RPM: Reads per million non-structural sRNA reads. The underlying data is the same as in panel **e**.

### TOFU-1:TOFU-2:SLFN-3/4 complexes process piRNA precursors

Next, we tested coIPs from BmN4 cells in which we co-expressed different combinations of TOFU-1, TOFU-2, SLFL-3 and SLFL-4 for piRNA processing activity. As substrate, we used a synthetic piRNA precursor oligonucleotide carrying an m7-G cap, which was radioactively labelled at its 3’-end with ^32^P for detection (**Fig.3a**). Processing activity was analyzed on a denaturing polyacrylamide gel system, alongside a synthetic RNA representing the expected processing product. This yielded processing activity, but only when both TOFU-1 and TOFU-2, as well as either SLFL-3 or SLFL-4, were present (**Fig.3b**). Introduction of an E216A mutation into TOFU-2 completely blocked this reaction (**Fig.3b**). Mammalian SLFN nucleases require divalent cations for cleavage activity^34,41^. Likewise, precursor processing was inhibited by EDTA, and was supported by divalent cations like Mg^2+^, Mn^2+^ or Ca^2+^ (at high concentrations), but not by Zn^2+^ (**Suppl. Fig.6a,b**).

To exclude the possibility that any BmN4-derived factors were responsible for the cleavage reaction, we also expressed both active and inactive (E216A) minimal versions of the TOFU-1-:TOFU-2:SLFL-3 complex recombinantly in *E. coli* (**Suppl. Fig.5f, g**). This minimal complex was active in precursor cleavage assays, while the E216A mutant was not (**Fig.3c**).

Mature piRNAs carry a monophosphate at their 5’ ends, which would be consistent with the cleavage product of a metal-dependent nuclease^34,42^. Successful ligation of the cleavage product to a synthetic RNA oligonucleotide with hydroxyl groups at both 5’ and 3’ ends confirmed the presence of a 5’-phosphate on the reaction product of the TOFU-1:TOFU-2:SLFL-3 nuclease (**Fig.3d**). Based on these results, we conclude that a complex of TOFU-1, TOFU-2 and either SLFL-3 or SLFL-4 constitutes the enzyme that processes the 5’-end of piRNA precursors in *C. elegans*. We name this complex PUCH for <u>P</u>recursor of 21<u>U</u> RNA 5’-end <u>C</u>leavage <u>H</u>oloenzyme.

### PUCH is a cap- and sequence-specific ribonuclease

We probed two key piRNA precursor properties for their relevance to processing. First, piRNA precursors are characterized by a 5’-m7-G Cap^24^. To examine whether the Cap-structure is essential for PUCH activity, we incubated full-length PUCH (isolated by TOFU-2 IP from BmN4 cell extracts) with a precursor having a 5’-phosphate (P) instead of a 5’-m7-G Cap. This experiment revealed that 5’-P precursor RNA was not processed, in contrast to the capped control substrate (**Fig.3e**). A second piRNA-precursor characteristic in *C. elegans* is the presence of a uracil at position three (U_3_)^24^. This corresponds to the most 5’ nucleotide in mature piRNAs, which display an extreme 5’U bias^23^. We tested whether PUCH could process a precursor substrate containing a cytosine at position three (AAC precursor) and found that PUCH did not cleave the AAC precursor at detectable levels (**Fig.3e**). Similar results for Cap- and U_3_-dependence were obtained with recombinant minimal PUCH complex that only contained the three SLFN domains of TOFU-1, TOFU-2 and SLFL-3 (**Fig.3c**), indicating that these features are recognized directly by the SLFN domains. We conclude that PUCH is a novel type of Cap- and sequence-specific ribonuclease.

### PUCH can cleave PETISCO-bound precursors

*In vivo*, piRNA precursors are bound by PETISCO^4,28^, and this enhances piRNA biogenesis. Yet, based on the results described thus far, PUCH does not require PETISCO for activity *in vitro*. PETISCO’s main role may therefore be to stabilize precursors *in vivo*, and not to promote PUCH activity. To genetically probe the relationship between PETISCO and PUCH, we asked how the loss of PETISCO function affects precursor accumulation in *tofu-2(e216a)* mutants. To this end, we sequenced small RNAs from a strain carrying the *tofu-2(e216)* allele and lacking the piRNA-specific PETISCO adapter protein PID-1^43^. In *tofu-2(e216a);pid-1(xf35)* double mutants precursor accumulation was reduced (**Fig.4a**), consistent with the idea that PETISCO stabilizes piRNA precursors to allow their processing by PUCH. Mature piRNAs were completely absent, as in *tofu-2(e216a)* single mutants (**Fig.4b**).

These results also imply that PUCH can process piRNA precursors while they are bound by PETISCO. To test this directly, we first incubated ^32^P-labelled precursors with purified PETISCO and tested binding in an electromobility-shift-assay (EMSA). We observed that the substrate was indeed bound by PETISCO, resulting in most of the complex not being able to enter the gel, most likely due to the large size of PETISCO (Octameric complex of 240 kDa^28^). The presence of a 5’-m7G-Cap on the precursor enhanced RNA binding by PETISCO (**Suppl. Fig.6c**). Next, we incubated PETISCO-precursor complexes with full-length immunopurified PUCH and analyzed cleavage products in a time-series, which revealed that cleavage is not prevented by the presence of PETISCO (**Fig.4c**). We conclude that PUCH can cleave piRNA precursors, also in presence of PETISCO.

### PUCH-PETISCO interaction affects precursor processing *in vivo*

If PUCH cleaves PETISCO bound precursors, interactions between the two complexes may be expected. However, multiple IP-MS experiments, including coIP of TOFU-2(E216A), did not reveal interactions between PUCH subunits and PETISCO (**Suppl. Fig.1d; Table S1; Table S2**). Reasoning that the presumed interaction may be too transient to be detected in *C. elegans* extracts, we systematically tested interactions between recombinant proteins in pull-down assays. This revealed an interaction between TOFU-1 and the PETISCO complex (**Fig.5a**). Using a combination of pulldown and size exclusion chromatography experiments, we narrowed down the interaction to a short segment of TOFU-1 (residues 82-113, TOFU-1^pep^), just N-terminal to the SLFN domain, and to the extended Tudor (eTudor) domain of the PETISCO subunit TOFU-6 (TOFU-6^eTUDOR^) (**Fig.5b**; **Suppl. Fig.7a-e**). Quantitative analysis with the minimal TOFU-1^pep^ using isothermal titration calorimetry revealed a *K*_d_ of ∼20 μM (**Suppl. Fig.7f**). We determined the crystal structure of the TOFU-6^eTUDOR^:TOFU-1^pep^ complex at 1.7 Å (**Fig.5c; TableS3**). TOFU-1^pep^ does not bind TOFU-6^eTUDOR^ at the canonical, dimethyl-arginine-binding aromatic cage of the TUDOR domain^44,45^, but on the surface of the staphylococcal nuclease-like domain (SN-like domain) of the eTudor domain (**Fig.5c; Suppl. Fig.8a,b**). This region has thus so far not been described to mediate protein-protein interactions. Based on the interaction interface, we designed mutations in both TOFU-1^pep^ and TOFU-6^eTUDOR^ that should disrupt their interaction and tested these using pull-downs and size exclusion chromatography (**Suppl. Fig.9a-e**). While mutations on only one of the partners (especially TOFU-1^pep^) weakened the interaction, mutation of both partners fully disrupted the interaction. We then tested the same mutations *in vivo*, using Crispr-Cas9 mediated mutagenesis of the endogenous loci, and sequenced piRNAs and their precursors from both single and double mutants. This revealed a reduction of mature piRNAs, as well as an accumulation of precursors, especially in the double mutants (**Fig.5d-f**). We conclude that piRNA accumulation *in vivo* is stimulated by the interaction between PETISCO and PUCH.

## Discussion

The identification of PUCH completes the piRNA biogenesis toolkit of *C. elegans*. At the sequence level PUCH is unrelated to Zuc, the enzyme that initiates piRNA biogenesis in mammals and flies. Yet, both enzymes perform a similar reaction: they both cleave piRNA precursors at a specified distance from the 5’-end of the precursor. While Zuc depends on Piwi proteins binding precursor 5’-ends^11,13,14^, PUCH depends on a 5’-m7G-Cap, which is likely bound by PUCH itself. A second commonality between Zuc and PUCH is the requirement of a uracil downstream of the cleaved phosphodiester bond. While for Zuc this is a rather weak requirement^11^, for PUCH this is a prerequisite for cleavage. This imposes a strong selection on potential novel sequences that may evolve towards piRNA precursors. A third similarity between the enzymes is that both contain a transmembrane helix. In case of PUCH, the transmembrane helix is located at the very C-terminus of SLFL-3/4. Even if we did not test the functional relevance of this domain in PUCH yet, Zuc is bound to the mitochondrial outer membrane via its N-terminal transmembrane helix^38,39^. The mere fact that both Zuc and PUCH share this feature suggests convergent evolution of membrane proximity of these enzymes, implying that recruitment to a membrane is important for piRNA 5’-end processing *in vivo*. Further experimentation will be required to resolve the role of membrane binding for Zuc and PUCH activity.

PUCH defines a novel type of ribonuclease, consisting of three subunits, each with one SLFN-like domain. How do our findings relate to SLFN-related proteins in other species? There is a variety of mammalian proteins that contain SLFN-folds. The *Slfn* gene cluster in mice has been described as an immunity locus, displaying high rates of sequence divergence^46^. Interestingly, a parental incompatibility syndrome, Dysdiadochokinesia (DDK) syndrome, has been linked to specific haplotypes of the *Slfn* gene cluster^46^. Given that enzymatic activity of PUCH requires association of three different SLFN-domain containing subunits, one can hypothesize that in mice complexes between distinct paternal and maternal Slfn proteins may form active enzymes, whose activity, or lack thereof, may trigger embryonic lethality. Another study in mice showed that a transposon encoded non-coding RNA inhibits *Slfn* gene expression and thus prevents over-activity of the innate immune system in response to virus infection^47^. Moreover, in humans links between immunity and SLFN proteins are known. For instance, SLFN11 restrains translation of viral proteins during HIV infection by cleaving specific tRNAs^48^. Interestingly, SLFN11 is a protein with multiple activities. SLFN11 binds single-stranded DNA, and it has been shown to also interfere with replication of certain DNA viruses and to be recruited to stalled replication forks^41^. Furthermore, members of the Orthopoxvirus family, such as the monkeypox virus, contain a virulence factor that carries a single SLFN domain^49,50^. Even though the relevance of this specific domain for virulence has not been assessed, a role in host-pathogen interaction control seems likely. Finally, a SLFN-related fold, the Smr domain, has been shown to act as a nuclease in RNA quality control mechanisms, and this function can be traced back to the last universal common ancestor^51^.

Overall, these activities, including the role we identify in piRNA biogenesis, point to a deeply conserved role for Slfn-like domains in immunity- and stress-related mechanisms. Our results show that SLFN domains can form multimeric complexes and that multimerization can unveil highly specific nucleolytic activities. It is conceivable that combinations between proteins with SLFN-related folds may enable uncharacterized activities that help organisms fighting off infectious nucleic acids.

## Supporting information

mass spec data 1

mass spec data 2

primers, strains etc

structure data

## Acknowledgements

We thank all members of the Ketting and Falk labs for fruitful discussions. This work was funded by the Deutsche Forschungsgemeinschaft (DFG, German Research Foundation) – Project-ID 439669440 – TRR 319. AWB was supported by the Peter und Traudl Engelhorn Foundation. Support by the IMB Genomics Core Facility and the use of its NextSeq500 (funded by the DFG – INST 247/870-1 FUGG) is gratefully acknowledged. Jan Schreier, Kay Holleis and Laurenz Miksch are acknowledged for their contribution to the early stages of this project. We also thank the beamline scientists from the European Synchrotron Radiation Facility (ESRF) beamline ID30A-3 (Grenoble, France) for excellent support with data collection.

## Author contributions

N.P. designed, executed, and analyzed the genetic experiments as well as the RNA binding and cleavage assays.

A.W.B. set up the experiments to produce PUCH complex from BmN4 cells, co-designed the cleavage assays and assisted in data analysis and interpretation.

R.L. Generated expression constructs, purified proteins for biochemical experiments and crystallization trials, performed protein interaction experiments.

S.H. generated *C. elegans* strains through genome editing.

E.N. and F.B. prepared samples for and analyzed them with mass spectrometry and analyzed the results.

T.F. purified proteins, designed and performed protein interaction experiments.

E.K. performed computational analysis of the small RNA data sets and visualized the results.

S.F. performed and interpreted AlphaFold predictions, ITC experiments and all crystallography related work.

R.F.K. assisted in data analysis and interpretation.

The study was conceived and designed by S.F. and R.F.K.

R.F.K., S.F., N.P. and A.W.B. contributed to writing the manuscript and making the figures, with input from all authors.

## Materials and correspondence

Sebastian Falk and René F. Ketting can be approached for materials and correspondence.

## Methods

### Worm culture

*C. elegans* strains were cultured on OP50 plates according to standard laboratory conditions^52^. For IP/MS experiment worms were grown on high-density egg OP50 plates^53^ and transferred to the standard OP50 plates for the last generation. The Bristol N2 strain was used as a reference wild-type strain. Used strains are listed in Table S4.

### CRISPR/CAS9 mediated genome editing

All protospacers were designed using CRISPOR (http://crispor.tefor.net) and afterwards confirmed with Integrated DNA Technologies “CRISPR-Cas9 guide RNA design checker”. Protospacers were cloned to the pRK2412 by SLIM (site directed ligase-independent mutagenesis). The Bristol N2 strain was used for microinjections, unless stated otherwise. ssDNA oligonucleotides (IDT) were utilized as repair template. Each of the repair templates has 35 nucleotides long homology arms. The injection mix contained 50ng/µl guide RNA coding plasmid for the gene of interest; 50ng/µl of plasmid, harboring CAS9 and *dpy-10* (cn64) or *unc-58(e665)* co-conversion guide RNA^54^; 750nM of ssDNA oligonucleotide (repair template for gene of interest) and 750nM of co-conversion ssDNA oligonucleotide. Used protospacers and repair templates are listed in the Table S4.

### Crosses with piRNA sensor

RFK1059 (*tofu-2*[E216A]) and *slfl-3(xf248)* mutant hermaphrodite worms were crossed with males of the RFK1246 strain, which carries a *mut-7* deletion as well as the piRNA sensor^22^. Worms, carrying piRNA sensor and *tofu-2[E216]* or *slfl-3(xf248)* mutation, and wild type for *mut-7* were selected by genotyping. Genotyping primers can be found in Table S4.

### Microscopy

Images of piRNA sensor carrying strains were obtained using a Leica DM6000B. Young adults and adult worms were washed in a drop of M9 (22mM KH_2_PO_4_, 42mM Na_2_HPO_4_, 85mM NaCl, 1mM MgSO_4_) and immobilized with 30mM sodium azide in M9 buffer. Imaging of Bm4 cells were done using Leica TCS SP5. Images were processed using ImageJ and Adobe Illustrator.

### Mass spectrometry

#### Worm pellet preparation

All IP/MS experiments were performed in quadruplicates. Worms, grown on the OP50 plates were bleached (2% NaClO, 666mM NaOH) into high-density egg plates, grown until gravid adult stage and bleached again. The embryos were left to hatch in M9 buffer (22mM KH_2_PO_4_, 42mM Na_2_HPO_4_, 85mM NaCl, 1mM MgSO_4_), L1 stage worms were seeded on standard OP50 plates and harvested at young adult stage. Worms were washed three times with M9 buffer and one time with cold sterile water. 200µl worm aliquots were pelleted and frozen in liquid nitrogen and stored at –80°C.

#### Lysis preparation

200 µl of synchronized young adult worms were thawed on ice and resuspended in 250 µl of 2x Lysis Buffer (50mM Tris HCl pH7.5, 300 mM NaCl, 3mM MgCl_2_, 2mM DTT, 2mM Triton X100, 2x cOmplete Mini, EDTA-free, Roche, 11836170001) and 50µl of sterile water. The Bioruptor Plus (Diagenode) sonicator was used to lyse worms (10 cycles 30/30 seconds, high energy, 4°C). After pelleting, the supernatant was accurately removed without the lipid phase. Finally, the protein concentration of the lysate was determined using Pierce BCA Protein Assay Kit (ThermoFisher Scientific, 23225).

#### Immunuoprecipitation

For anti-HA IPs, 550µL of worm lysate containing 0,75mg protein was resuspended in a final volume of 550µL of 1x Lysis Buffer. Anti-HA IPs were performed with 2µg of in-house made anti-HA antibodies (mouse, clone 12CA5). The lysate was incubated with the antibodies for two hours at 4°C. For each sample, 30µl of protein G magnetic beads (Dynabeads, Invitrogen) were washed three times in washing buffer (25mM Tris HCl pH7.5, 150mM NaCl, 1.5mM MgCl_2_, 1mM DTT, 1mM Triton X100, cOmplete Mini, EDTA-free, Roche, 11836170001). Subsequently, equilibrated beads were added to the lysis and incubated for an additional hour at 4°C by end-over-end rotation. Finally, beads were washed 6 times with Wash Buffer, resuspended in 2xNuPAGE LDS Sample Buffer (containing 200mM DTT) and boiled for 15min at 95°C.

To identify of TOFU-2::HA and TOFU-2::HA(E216A), samples were separated on a 4%–12% NOVEX NuPAGE gradient SDS gel (Thermo) for 10 min at 180 V in 1× MES buffer (Thermo). Proteins were fixated and stained with Coomassie G250 Brilliant Blue (Carl Roth). The gel lanes were cut, minced into pieces, and transferred to an Eppendorf tube. Gel pieces were destained with a 50% ethanol/ 50mM ammonium bicarbonate (ABC) solution. Proteins were reduced in 10mM DTT (Sigma-Aldrich) for 1h at 56°C and then alkylated with 5mM iodoacetamide (Sigma-Aldrich) for 45 min at room temperature. Proteins were digested with trypsin (Sigma) overnight at 37°C. Peptides were extracted from the gel by two incubations with 30% ABC/acetonitrile and three subsequent incubations with pure acetonitrile. The acetonitrile was subsequently evaporated in a concentrator (Eppendorf) and loaded on StageTips^55^ for desalting and storage.

For mass spectrometric analysis, peptides were separated on a 20-cm self-packed column with 75µm inner diameter filled with ReproSil-Pur 120 C18-AQ (Dr.Maisch GmbH) mounted to an EASY HPLC 1000 (Thermo Fisher) and sprayed online into an Q Exactive Plus mass spectrometer (Thermo Fisher). We used a 94-min gradient from 2 to 40% acetonitrile in 0.1% formic acid at a flow of 225nl/min. The mass spectrometer was operated with a top 10 MS/MS data-dependent acquisition scheme per MS full scan. Mass spectrometry raw data were searched using the Andromeda search engine^56^ integrated into MaxQuant suite 1.6.5.0^57^ using the UniProt *C. elegans* database (August 2014; 27,814 entries). In both analyses, carbamidomethylation at cysteine was set as fixed modification, while methionine oxidation and protein N-acetylation were considered as variable modifications. Match-between-run option was activated. Prior to bioinformatic analysis, reverse hits, proteins only identified by site, protein groups based on one unique peptide, and known contaminants were removed.

For the further bioinformatic analysis, the LFQ values were log2-transformed and the median across the replicates was calculated. This enrichment was plotted against the – log 10-transformed P value (Welch’s t-test) using the ggplot2 package in the R environment.

### Western blot from the worm lysis

Worms were grown and lysed as described for the Mass spectrometry section. Lysis of both RFK1269 and RFK1280 worms, containing 15µg of protein, were mixed with 2x gel loading buffer (2x Novex NuPage LDS sample buffer (Invitrogen), supplemented with 200mM DTT) and were heated at 95°C for 10min prior to resolving on a 4-12% Bis-Tris NuPage NOVEX gradient gel (Invitrogen) in 1x Novex NuPAGE MOPS SDS Running Buffer (Invitrogen) at 150 V. Separated proteins were transferred to nitrocellulose membrane (Amersham) 1h at 120V using 1x NuPAGE Transfer Buffer (Invitrogen) supplemented with 10% methanol. The membrane was incubated for 30min in 1x PBS-Tween (0.05%) supplemented with 5% skim milk, cleaved and incubated overnight with primary antibodies diluted in PBS-Tween (1:1,000 monoclonal anti-HA (12CA5, in-house); 1:1,000 anti-h3 (H0164, Sigma) rabbit polyclonal antibodies. Subsequently, the membrane was washed 5 times for 5min in PBS-Tween, prior to incubation with the secondary antibody, using 1:10,000 horse anti-mouse HRP-linked antibody (#7076, Cell Signaling) and goat anti-rabbit HRP-linked antibodies (#7074, Cell Signaling) and imaged using SuperSignal™ West Pico Plus (TermoFischer) kit.

### RNA isolation and RNA sequencing

Worms were grown at 20°C, synchronized by bleaching (2% NaClO, 666mM NaOH) and were left to hatch overnight in M9 buffer. Next, L1-stage worms were seeded onto OP50 plates and harvested as young adults. For RNA extraction 500 µL of TRISOL LS (ThermoFisher Scientific, 10296-028) was added to the 50µL worm aliquot, and five cycles of freezing in liquid nitrogen/thawing in the 37°C water bath were performed. Samples were centrifuged for 5min at 21x*g* at RT, and supernatant was collected. 1 volume of 100% EtOH was added to the supernatant, before proceeding with the RNA extraction using the Direct-zol™ RNA MicroPrep (Zymo) kit. RNA was eluted into 13µl of Nuclease Free water (Ambion^®^ Invitrogen™) and each sample was divided into two aliquots for piRNA-precursor and mature piRNA library preparation.

#### CIP/RppH treatment and library preparation (for precursors)

CIP treatment on 1,5µg of isolated RNA was performed in rCutSmart™ Buffer (B6004S) using 3µL of Quick CIP (M0525L) in a 40µL reaction. The reaction was incubated at 37°C for 20min, followed by heat inactivation for 2min at 80°C. The CIP-treated RNA was subjected to another round of purification using the Direct-zol™ RNA MicroPrep (Zymo) kit. RppH (NEB) treatment was performed with a starting amount of 500ng.

#### Library preparation, sequencing and analysis

NGS library prep was performed with NEXTflex Small RNA-Seq Kit V3 following Step A to Step G of Bioo Scientific’s standard protocol. Amplified libraries were purified by running an 8% TBE gel and size-selected for 15-40 nt. Libraries were profiled in a High Sensitivity DNA Chip on a 2100 Bioanalyzer (Agilent technologies), quantified using the Qubit dsDNA HS Assay Kit, in a Qubit 2.0 Fluorometer (Life technologies) and sequenced on Illumina NextSeq 500/550.

The raw sequence reads in FastQ format were cleaned from adapter sequences and size-selected for 18-35 nt inserts (plus 8 random adapter bases) using cutadapt v.4.0 (http://cutadapt.readthedocs.org) with parameters “-a TGGAATTCTCGGGTGCCAAGG -m 26 -M 43”. Data quality was assessed with FastQC v.0.11.9 (https://github.com/s-andrews/FastQC) and MultiQC v.1.9 (https://multiqc.info/). Read alignment to the *C. elegans* genome (Ensembl WBcel235/ce11 assembly) with concomitant trimming of the 8 random bases was performed using Bowtie v.1.3.1 (http://bowtie-bio.sourceforge.net) with parameters “-v 1 -M 1 -y --best --strata --trim5 4 --trim3 4 -S” and the SAM alignment files were converted into sorted BAM files using Samtools v.1.10 (http://www.htslib.org). *C. elegans* WBcel235/ce11 gene annotation in GTF format was downloaded from Ensembl release 96 (ftp://ftp.ensembl.org/pub/). Aligned reads were assigned to small RNA loci and classes using Samtools, GNU Awk and Subread featureCounts v.1.6.2 (http://bioinf.wehi.edu.au/featureCounts/). Structural reads aligned in sense orientation to rRNA, tRNA, snRNA and snoRNA loci were excluded from further analysis. Mature piRNAs were stringently defined as reads of length 21nt starting with T and fully overlapping with annotated piRNA (21ur) genes in sense orientation. Because 21ur gene annotation corresponds to mature piRNA sequences, piRNA precursors were stringently defined as reads of length 23-35 nt starting 2 bp upstream of the annotated 5’-end of (mature) piRNAs in sense orientation. The relative abundance of mature and precursor piRNAs was normalized to the number of non-structural 18-35 nt reads in each sample. Coverage tracks of aligned reads overlapping in sense with piRNA genes were produced using Bedtools v.2.27.1 (http://bedtools.readthedocs.io) and kentUtils v.385 (https://github.com/ucscGenomeBrowser/kent). The tracks were normalized based on all non-structural reads in each sample and visualized on the IGV genome browser v.2.15.4 (https://igv.org/).

### 3’ RNA radioactive labeling

3’-end labeling of substrate RNA (see Table S4 for sequence) was performed in a 25µL reaction containing 2.5µL f DMSO, 2.5µL of T4 ligase buffer (NEB), 1µL of T4 ligase (NEB), 2.5µL 10mM ATP (NEB), 1µL of synthetic RNA precursor (5pmol/µL). The reaction was mixed and and 2.5µL of [5’-^32^P]pCp (SCP-111, Hartmann analytic) was added before overnight incubation at 16°C. Finally, the labeled RNA was purified using G25 columns (Cytiva) according to the manufacturer’s protocol. The 3’-end labeled synthetic RNA precursor was used for *in vitro* cleavage assays and in EMSAs.

### 5’ RNA radioactive labeling

5pmol synthetic RNA oligonucleotide was labeled with ATP, [γ-^32^P] (PerkinElmer) using T4 PNK(NEB), according to the manufacturer’s protocol. The sequences of the RNA substrates can be found in Table S4.

### Plasmids

Full-length CeTOFU-2 was amplified from N2 cDNA and was inserted by restriction-based cloning into the pBEMBL vector (kind gift of R.Pillai) in which expression of a N-terminal eGFP tag is driven by the OpIE2 promoter. Likewise, CeTOFU-1 was inserted into a vector harboring a C-terminal 3xFLAG-mCherry cassette. CeSLFL3 and CeSLFL4 were inserted into a vector backbone containing an N-terminal HA tag. All primers, vector backbones and detailed cloning strategies can be found in Supplemental Materials.

### BmN4 cell culture and transfection

BmN4 cells were cultured at 27°C in IPL-41 insect medium (Gibco) supplemented with 10% FBS and 0.5% Pen-Strep (Gibco). 24 hours prior to transfection, ∼4x 10^6 cells were seeded in a 10-cm dish (using one 10-cm dish for each condition in the cleavage reaction). Cells were transfected with 10 µg of each plasmid DNA using XtremeGene HP (Roche) transfection reagent, according to the manufacturer’s instructions. 72h post transfection cells were harvested, washed twice in ice-cold PBS and pelleted by centrifugation at 500xg for 5min at 4°C.

### GFP-IP from BmN4 cells

Roughly 4 × 10^^6^ BmN4 cells were harvested from each 10-cm dish (see above), washed once in 5mL ice-cold PBS and once more in 1 mL ice-cold PBS. Subsequently, cells were pelleted by centrifugation for 5min at 500xg at 4°C and frozen at -80°C. Directly before use, BmN4 cell pellets were thawn on ice and lysed in 1 mL IP-150 Lysis Buffer (30mM Hepes [pH7.4], 150mM KOAc, 2mM Mg(OAc)_2_ and 0.1% Igepal freshly supplemented with EDTA-free protease inhibitor cocktail and 5mM DTT) for 1 hour by end-over-end rotation at 4°C. Cells were further lysed by passing the lysate ten times through a 20-gauge syringe needle followed by five passes through a 30-gauge needle. Cell debris was pelleted by centrifugation at 17,000xg for 20min at 4°C. Supernatant fractions were collected and subjected to GFP-IP using GFP-Trap beads (Chromotek). The GFP-Trap beads (15µL beads suspension per reaction) were washed 3 times in 1mL of IP-150 Lysis Buffer. Equilibrated beads were subsequently incubated with the BmN4 cell lysate and incubated overnight by end-over-end rotation at 4°C. The next day, immunoprecipitated complexes were washed five times using 1mL of IP-150 Lysis Buffer and were subsequently used for *in vitro* cleavage assays or for immunodetection using Western Blot analysis.

### Western Blot

Samples were prepared in 1x Novex NuPage LDS sample buffer (Invitrogen) supplemented with 100mM DTT and were heated at 95°C for 10min prior to resolving on a 4-12% Bis-Tris NuPage NOVEX gradient gel (Invitrogen) in 1x Novex NuPAGE MOPS SDS Running Buffer (Invitrogen) at 140V. Separated proteins were transferred to nitrocellulose membrane (Amersham) overnight at 20V using 1x NuPAGE Transfer Buffer (Invitrogen) supplemented with 10% methanol. The next day, the membrane was incubated for 1h in 1x PBS-Tween (0.05%) supplemented with 5% skim milk and incubated for 1 hour with primary antibodies diluted in PBS-Tween (1:1,000 monoclonal anti-Flag M2, F3165, Sigma-Aldrich; 1:1,000 monoclonal anti-GFP antibodies (B-2), Santa Cruz, sc-9996, K1115; 1:1,000 monoclonal anti-HA (12CA5, in-house); 1:1,000 anti-actin (A5060) rabbit monoclonal antibodies, Sigma. Subsequently, the membrane was washed 3 times for 5min in PBS-Tween, prior to incubation with the secondary antibody, using 1:10,000 IRDye 800CW Goat anti-mouse and IRDye 680LT Donkey anti-rabbit IgG (LI-COR) and imaged on an Odyssey CLx imaging system (LI-COR).

### *In vitro* cleavage assay

The PUCH complex used for *in vitro* cleavage assays was obtained in two different ways. The full-length PUCH complex was obtained from GFP-IPs using BmN4 cell lysates (see above), whereas the minimal catalytic complex (mini-PUCH) was purified from *E*.*coli*.

For the *in vitro* cleavage assays performed with IP material from BmN4 cells, beads were washed in the cleavage buffer (CB) containing 40mM Tris-HCl, pH 8.0, 20mM KCl, 11mM MgCl_2_ and 2mM DTT. Beads were subsequently resuspended in 10µL of CB and incubated with 0.2pmol of the labeled RNA substrate for 1h at room temperature.

For cleavage assays with mini-PUCH purified from *E*.*coli* 0.2pmol of labeled RNA substrate was incubated in 10µL CB buffer with 27nM mini-PUCH protein complex (final concentration) at 20°C for 30min.

The cleavage reaction was terminated by adding 1µL of 20mg/ml proteinase K. 1 volume of the 2xRNA Gel Loading Dye (Thermo Scientific™, R0641) was added and the RNA was resolved on a 15% TBE-UREA gel (Novex™) for 90min at 180V with 1xTBE as the running buffer.

#### Substrate specificity test of PUCH complex

Capped RNA oligonucleotides were labeled at the 3’end, 0.2pmol (1µL) of RNA per sample were used in the cleavage reaction. For reaction with IP material, to obtain 5’P-containing piRNA precursor oligonucleotide, 5’OH-piRNA precursor had been labeled on the 3’end as described above. After labeling, 5’P was created by T4 PNK treatment (NEB, M0201S), done according to the NEB T4 PNK protocol. For the reaction with mini-PUCH 5’OH-piRNA precursor oligonucleotide had been labeled on the 5’end as described above.

#### Analysis of divalent cations as cofactor of PUCH complex

For the metal assay, beads were washed with CB, but 100mM MgCl_2_ was replaced by ZnCl_2_, MnCl_2_ or CaCl_2_. Cleavage reaction was done with full-length PUCH obtained from GFP-IP material from BmN4 cell lysates.

#### Ligation of small RNA oligo to the cleavage product in order to prove formation of 5’P on the cleaved RNA precursor

2pmol of labeled RNA was incubated in 35µL of CB containing mini-PUCH (or mutated mini-PUCH) at a final concentration of 40nM and was incubated at 20°C for 1h. Afterwards 3 volumes of Trisol LS reagent (ThermoFisher Scientific, 10296-028) was added, and RNA was purified using Direct-zol™ RNA MicroPrep (Zymo) according to the manufacturer’s protocol. Next, the RNA was ligated to 10pmol of 5’OH-rGrUrCrUrGrUrUrUrArA-OH3’ oligonucleotide using T4 RNA ligase according to the manufacturer’s protocol. After 16h of incubation at 16°C, the reaction terminated by proteinase K and RNA was resolved on a 15% TBE-UREA gel (Novex™) for 90min at 180V with 1xTBE as the running buffer.

#### PUCH complex cleavage activity in presence of PETISCO

The assay has been done with PUCH complex, obtained from Bm4 cells. Per sample: 1µL of 3’end labeled piRNA precursor (0.2pmol) was incubated with five times access of PETISCO protein complex on ice for 1h in 10µL of CB buffer. PUCH-IP containing beads were resuspended in 10µL of RNA-PETISCO mix and incubated at 20°C.

### EMSA

0.2pmol of capped piRNA precursor, 5’P piRNA precursor and 5’OH-piRNA precursor were incubated with recombinant proteins of PETISCO complex, containing IFE-3, TOFU-6, ERH-2 and PID-3^28^ in a concentration range from 75pM to 1.44µM, in 10μL of binding buffer (20mM HEPES pH7.5, 150mM NaCl) for 1h at the room temperature. After the incubation, each sample was mixed with 15% Ficoll with bromophenol blue. Native 6% TBE gel was pre-run for 30min at 180V at room temperature in 1xTBE, and samples were resolved for 2h.

### Recombinant protein production in *E. coli*

PETISCO and its subunits (IFE-3, TOFU-6, PID-3, ERH-2) were purified and reconstituted as described in^28^. Using ligation-independent cloning, genes encoding TOFU-1, TOFU-2, and SLFL-3 were cloned into modified pET vectors. All proteins were produced as an N-terminal His-Tagged fusion protein with varying fusion tags that can be removed by the addition of 3C protease. Proteins or protein complexes were produced in the *E. coli* BL21(DE3) derivate strains in terrific broth medium. Briefly, cells were grown at 37°C, and when the culture reached an optical density (OD) at 600 nm of 2-3, the temperature was reduced to 18°C. After 2h at 18°C, 0.2mM IPTG was added to induce protein production for 12-16 h overnight.

### Co-expression pull-down assays

For interaction studies by the co-expression co-purification strategy, two plasmids containing the gene of interest and different antibiotic resistance markers were co-transformed into BL21(DE3) derivative strains to allow co-expression. 50 mL of cells were grown in TB medium shaking at 37°C, and when the culture reached an OD at 600 nm of 2-3, the temperature was reduced to 18°C. Protein production was induced after 2 h at 18°C, through the addition of 0.2 mM IPTG for 12-16 h overnight. Cells were harvested by centrifugation and the cell pellets were resuspended in 2 mL of lysis buffer (50 mM Sodium phosphate, 20 mM Tris/HCl, 250 mM NaCl, 10 mM Imidazole, 10% (v/v) glycerol, 0.05% (v/v) NP-40, 5 mM 2-mercaptoethanol pH 8.0) per gram of wet cell mass. Cells were lysed by ultrasonic disintegration, and insoluble material was removed by centrifugation at 21,000xg for 10 min at 4°C. For Streptactin pull-downs, 500 µL of supernatant was applied to 20 Strep-Tactin^®^XT resin (IBA Lifesciences); for MBP pull-downs, 500 µL supernatant was applied to 20 µL amylose resin (New England Biolabs) and incubated for two hours at 4°C. Subsequently, the resin was washed three times with 500 µL of lysis buffer. The proteins were eluted in 50 µL of lysis buffer supplemented with 10 mM maltose or 50 mM biotin in the case of amylose beads or Strep-Tactin^®^XT beads, respectively. Input material and eluates were analyzed by SDS–PAGE and Coomassie brilliant blue staining.

### Pull-down assays with purified proteins

To analyze protein interaction with purified proteins, appropriate protein mixtures (bait 10-20 µM, prey in 1.2-fold molar excess) were incubated in binding buffer containing 20 mM Tris/HCl (pH 7.5), 150 mM NaCl, 10% (v/v) glycerol, 0.05% (v/v) NP40, 1 mM DTT for 30min at 4°C. Subsequently, the indicated beads were added to the protein mixtures were then incubated with the indicated beads for 2 h on ice: Glutathione sepharose beads (Cube Biotech), Amylose sepharose beads (New England Biolabs)), and Strep-Tactin XT beads (IBA). Subsequently, the beads were washed three times with 200 μL mL binding buffer, and the retained material was eluted with 0.05 mL incubation buffer supplemented with 20 mM of reduced glutathione, 10 mM maltose, or 50 mM biotin. Input material and eluates were analyzed by SDS–PAGE and Coomassie brilliant blue staining.

### Purification of the trimeric mini PUCH

To reconstitute minimal PUCH, TOFU-1 (residues 160 to 373), TOFU-2 (residues 200-433) and SLFL-3 (residues 1-345) were co-expressed in BL21 (DE3). In case of inactive minimal PUCH, an inactive TOFU-2 mutant (residues 200-433, E216A) was used. TOFU-1 carried an N-terminal His10-MBP tag, TOFU-2 an N-terminal His10-MBP and a C-terminal StrepII tag, and SLFL-3 an N-terminal His6-GST tag. Cells were grown at 37°C, and when the culture reached an optical density (OD) at 600 nm of 2-3, the temperature was reduced to 18°C. After 2 h at 18°C, 0.2 mM IPTG was added to induce protein production for 12-16 h overnight. All purification steps were performed on ice or at 4°C. Cells were lysed by sonication in lysis buffer (50 mM Sodiumphosphate, 20 mM Tris/HCl, 500 mM NaCl, 20 mM imidazole, 10% (v/v) glycerol, and 5 mM 2-mercaptoethanol at pH 8.0). PUCH was purified by immobilized metal affinity chromatography (IMAC) using a 5 mL Ni2+-chelating HisTrap FF column (Cytiva). Proteins were eluted with lysis buffer supplemented with 500 mM imidazole and dialyzed overnight against 20 mM Tris/HCl, 150 mM NaCl, 10% (v/v) glycerol, and 5 mM 2-mercaptoethanol at pH 7.5. After dialysis, PUCH was subjected to heparin affinity chromatography on a 5 mL HiTrap Heparin HP (Cytiva) followed by size exclusion chromatography on a HiLoad Superdex 200 16/600 (Cytiva) in 20 mM Tris/HCl pH 7.5, 150 mM NaCl, 10% (v/v) glycerol, 2 mM DTT.

### Analytical size exclusion chromatography

Purified proteins were incubated alone or in different mixtures in concentrations between 20-40 µM (total volume of 50 µl) in size exclusion buffer (20 mM Tris/HCl pH 7.5, 150 mM NaCl, 2 mM DTT) as indicated in the figure legends. Samples were incubated for 1 h on ice to allow complex formation. Complex formation was assayed by comparing the elution volumes in SEC on a Superdex 200 Increase 3.2/300 (Cytiva). The SEC peak fractions were analyzed by SDS–PAGE and visualized by Coomassie brilliant blue staining.

### Isothermal titration calorimetry

ITC experiments to quantitatively analyze the interaction between TOFU-1 peptide (residues 82-113) and the TOFU-6 eTUDOR domain (residues 119-314) interaction were carried out using the PEAQ-ITC Isothermal titration calorimeter (Malvern). The TOFU-1 82-113 peptide does not contain Tyrosine or Tryptophane residues. To be able to determine the concentration precisely, we engineered a TOFU-1 peptide (TOFU-1^W-82-113^) that contains a Tryptophan residue at the N-terminus. Data processing and analysis was performed using the PEAQ-ITC software (Malvern). Before the measurements, the samples were dialyzed overnight simultaneously against 1 L of ITC buffer (20 mM Tris, 250 mM NaCl, 0.5 mM TCEP, pH 7.50). TOFU-1^W-82-113^ (the reactant) samples were concentrated to 45-48 µM and TOFU-6^eTUDOR^ (the injectant) to 400-450 µM. Titrations were carried out at 25°C with 2 µL of the injectant per injection added to 200 µL of reactant cell solution. The reported Kd and stoichiometry are the average of three experiments, and the reported experimental error is the standard deviation.

### TOFU-6^eTUDOR^ TOFU-1^pep^ crystallization

Purified TOFU-6^eTUDOR^ and TOFU-1^W-82-113^ were mixed with TOFU-1^W-82-113^ being in 1.5-fold molar excess and subjected to size exclusion chromatography on a HiLoad Supdersex S75 16/600 (Cytiva) equilibrated in 20 mM Tris/HCl, 150 mM NaCl, 2 mM DTT pH7.5. The complex containing fractions were concentrated to 10 mg/ml by ultrafiltration. Crystallizsation trials were performed at 4°C and 22°C at 8-10 mg/ml using a vapor diffusion set-up. Drops were setup using a mosquito^®^ Crystallization Robot (SPT Labtech) on 96-Well 2-Drop MRC Crystallization Plates (Swissci) by mixing the protein complex and crystallization solution in a 200:200 nl and 400 :200 nl ratio.

Small crystals grew at 4°C in various conditions of the Morpheus Screen^58^. Several rounds of microseed matrix screening yielded larger crystals. The best crystals grew in 0.2 M Na bromide, 0.1 M Bis Tris propane pH 7.5, 20% (w/v) PEG 3350 at 22°C in the PACT screen^59^. Crystals were soaked with a mother liquor supplemented with 20% (v/v) glycerol for cryoprotection and then frozen in liquid nitrogen.

### Data collection, structure determination and refinement

Data were collected at the ESRF (Grenoble, France) beamline ID30A-3 on September 26^th^ 2021; https://doi.esrf.fr/10.15151/ESRF-DC-1033968485.

Data were processed with autoPROC^60^ using XDS^61^ and AIMLESS^62^. Phases were determined by molecular replacement using the AlphaFold model of the *C. elegans* TOFU-6 eTUDOR domain (residues 120-314) (https://alphafold.ebi.ac.uk/entry/Q09293). Molecular replacement was performed with Phaser^63^ within Phenix^64^. The model was processed with Phenix (process predicted model) to translate the pLDDT values to B factors and to remove flexible regions. Following molecular replacement, the model was automatically built with Buccaneer^65^, manually completed with COOT^66^ and refined using phenix.refine^67^. Model quality was assessed using molprobity^68^ and PDB-REDO^69^. The refined model has a clashscore of 5.06 and 98.54% of the residues fall into Ramachandran favored and 1.46% into Ramachandran allowed regions. Data collection and refinement statistics are listed in Supplemental Table S3. Molecular graphics of the structures were prepared using UCSF ChimeraX^70^. Coordinates and structure factors have been deposited in the Protein Data Bank with accession codes PDB ID 8BY5.

### Protein complex structure prediction with AlphaFold2 multimer

The prediction of protein complex structures was performed using AlphaFold v2.1.0 on the Colab notebook (ColabFold)^71^: https://colab.research.google.com/github/sokrypton/ColabFold/blob/main/AlphaFold2.ipynb The following settings were used: template_mode (none), msa_mode (MMSeq2 (UniRef+Environmental), pair_mode (unpaired + paired), model_type (AlphaFold2-multimer-v2). In case of the tetrameric PUCH 3 recycles were used, for the trimeric PUCH, 48 recycles were used. Protein sequences were obtained from Uniprot and for initial complex predictions, full-length sequences from all four proteins (TOFU-1, TOFU-2, SLFL-3, and SLFL-4) were used. The predicted models and the predicted alignment error (PAE) score were visualized and analyzed using ChimeraX^70^. Predicted complexes either contained SLFL-3 or SLFL-4. As SLFL-3 and SLFL-4 are paralogs that are 90% identical and 93% similar on the protein sequence level, we focused on predictions of the trimeric PUCH containing TOFU-1, TOFU-2 and SLFL-3. For the prediction of the core PUCH the following residue boundaries were used. TOFU-1 residues 156-373, encompassing the SLFN domain with an N-terminal extension. TOFU-2 residues 200-433, encompassing the SLFN domain and two C-terminal alpha helices. SLFL-3 residues 103-300, encompassing the SLFN domain.

## Data availability

Sequencing data is available at NCBI’s Sequence Read Archive under accession number PRJNA925182 (https://dataview.ncbi.nlm.nih.gov/object/PRJNA925182?reviewer=951lavsn8a0umpj5j17m8bk738). The mass spectrometry proteomics data have been deposited to the ProteomeXchange Consortium via the PRIDE^72^ partner repository with the dataset identifier PXD039502.

Username

Password: T2C6anBX

Coordinates and structure factors of the TOFU-6^eTUDOR^ TOFU-1^pep^ complex structure have been deposited in the Protein Data Bank with accession codes PDB ID 8BY5.

## Code availability

Codes can be obtained upon reasonable request from the corresponding authors.

---

**Fig. S1.**
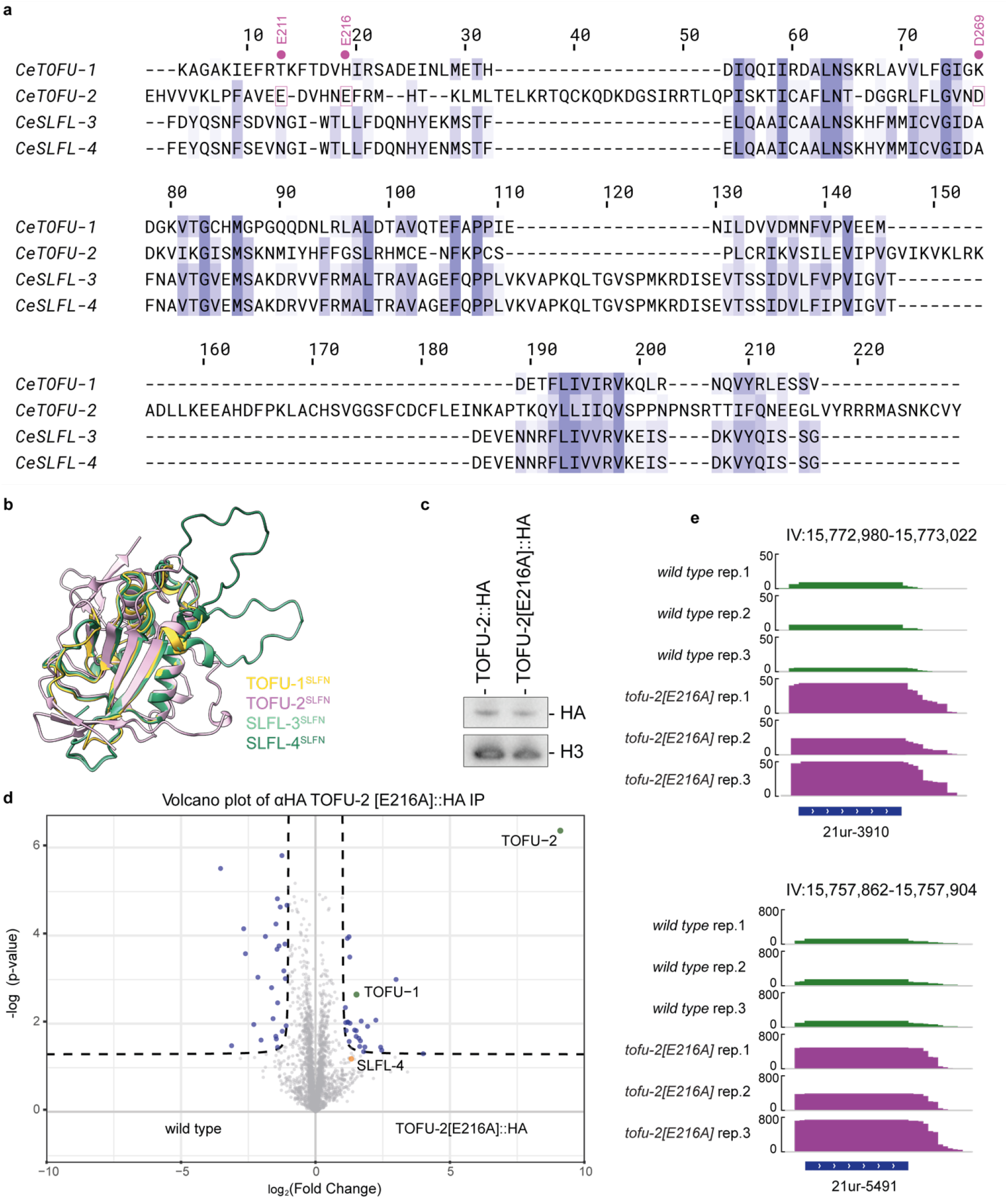
Mutation in the catalytic center of TOFU-2 does not affect protein stability and interaction with TOFU-1. **a**, Structure-based sequence alignment of the SLFN domains from TOFU-1, TOFU-2, SLFL-3 and SLFL-4. The acidic residues from the active site of TOFU-2 are highlighted with purple boxes and the residue number is indicated on the top. **b**, Superposition of the AlphaFold predicted SLFN domains fromTOFU-1, TOFU-2, SLFL-3 and SLFL-4. TOFU-1 is shown in yellow, TOFU-2 in purple and the paralogs SLFL-3/4 in different shades of green. **c**, Extracts of young adult worms with genotype *tofu-2::HA* and *tofu-2[E216A]::HA*, were separated on SDS-PAGE. Western blot was probed using anti-HA and anti-H3 antibodies, followed by visualization with HRP. **d**, Volcano plot representing label-free proteomic quantification of TOFU-2[E216A]::HA (n=4) and wild type (n=4, WT) immunoprecipitations from young adult extracts. The X-axis represents the median fold enrichment of individual proteins in wild type (WT) versus the TOFU-2::HA mutant strain. The Y-axis indicates −log10(P-value) calculated using Welch t-test. Dashed lines represent enrichment thresholds at p-value = 0.05 and fold change > 2, c = 0.05. Each dot represents an enriched (blue/grey) or quantified (orange) proteins. **e**, Genome browser tracks of two individual piRNA loci, displaying normalized read coverage in piRNA precursor libraries. Top three tracks (green) are from a wild-type background, the bottom three tracks (purple) are from a *tofu-2[E216A]* mutant background.

**Fig. S2.**
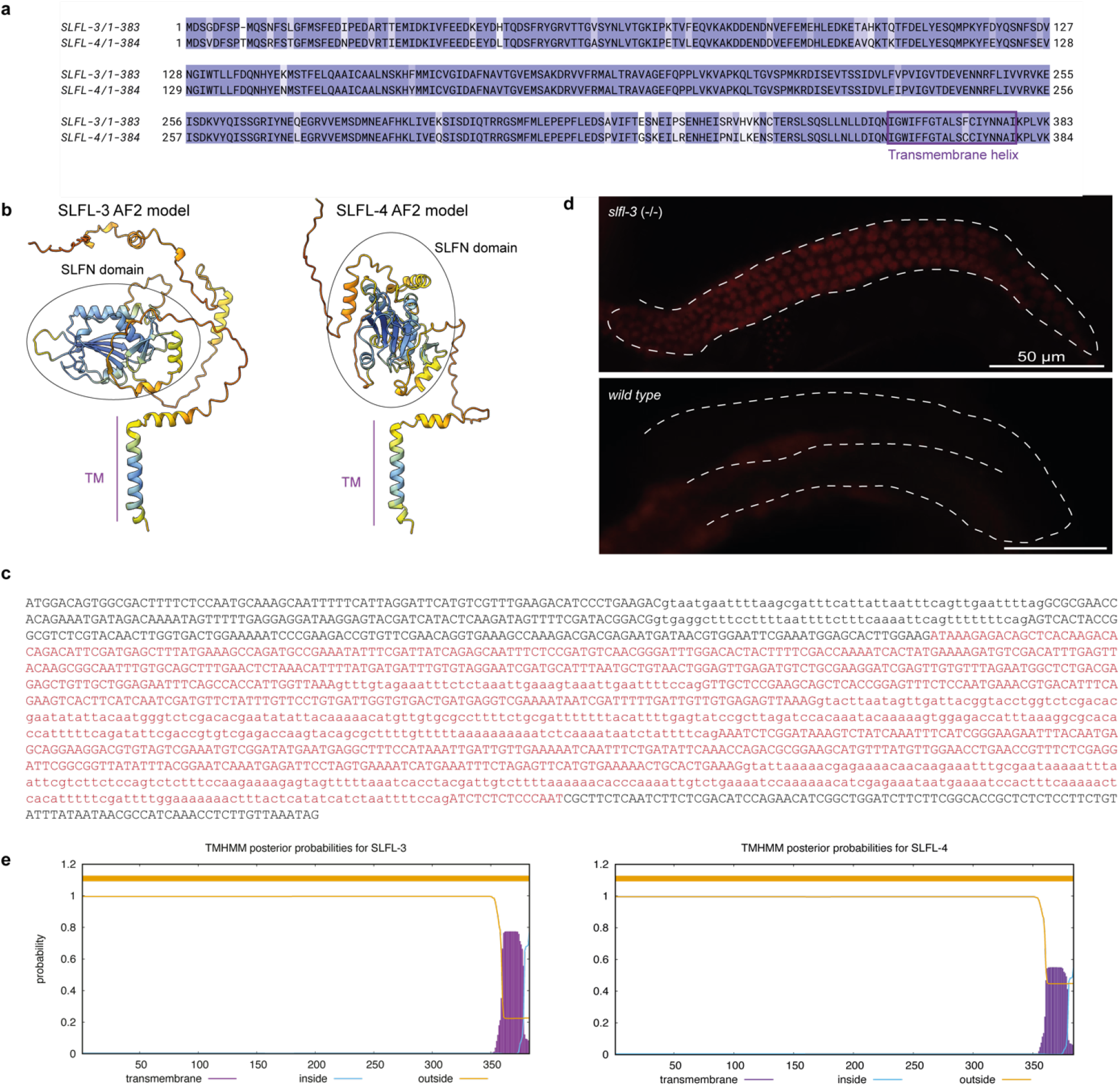
SLFL-3 and SLFL-4 are transmembrane proteins involved in piRNA biogenesis in *C. elegans*. **a**, Sequence alignment of SLFL-3 and SLFL-4. The predicted C-terminal transmembrane helix is highlighted with a box. **b**, AlphaFold2 predicted structures of SLFL-3/4 shown as cartoon and colored by pLDDT score, which reports on the model confidence. Dark blue indicates very high, light blue confident, yellow low and orange very low model confidence. **c**, Sequence of *slfl-3(xf248)* allele. Deleted sequence is marked in red. **d**, Widefield fluorescent microscopy of adult hermaphrodites carrying the mCherry::H2B-piRNA sensor in two genetic backgrounds: *slfl-3(xf248)* on top and wild type at the bottom. The germlines are outlined by a white dashed line. Scale bar – 50 µm. **e**, Prediction of transmembrane helices in SLFL-3 and SLFL-4 using TMHMM - 2.0.

**Fig. S3.**
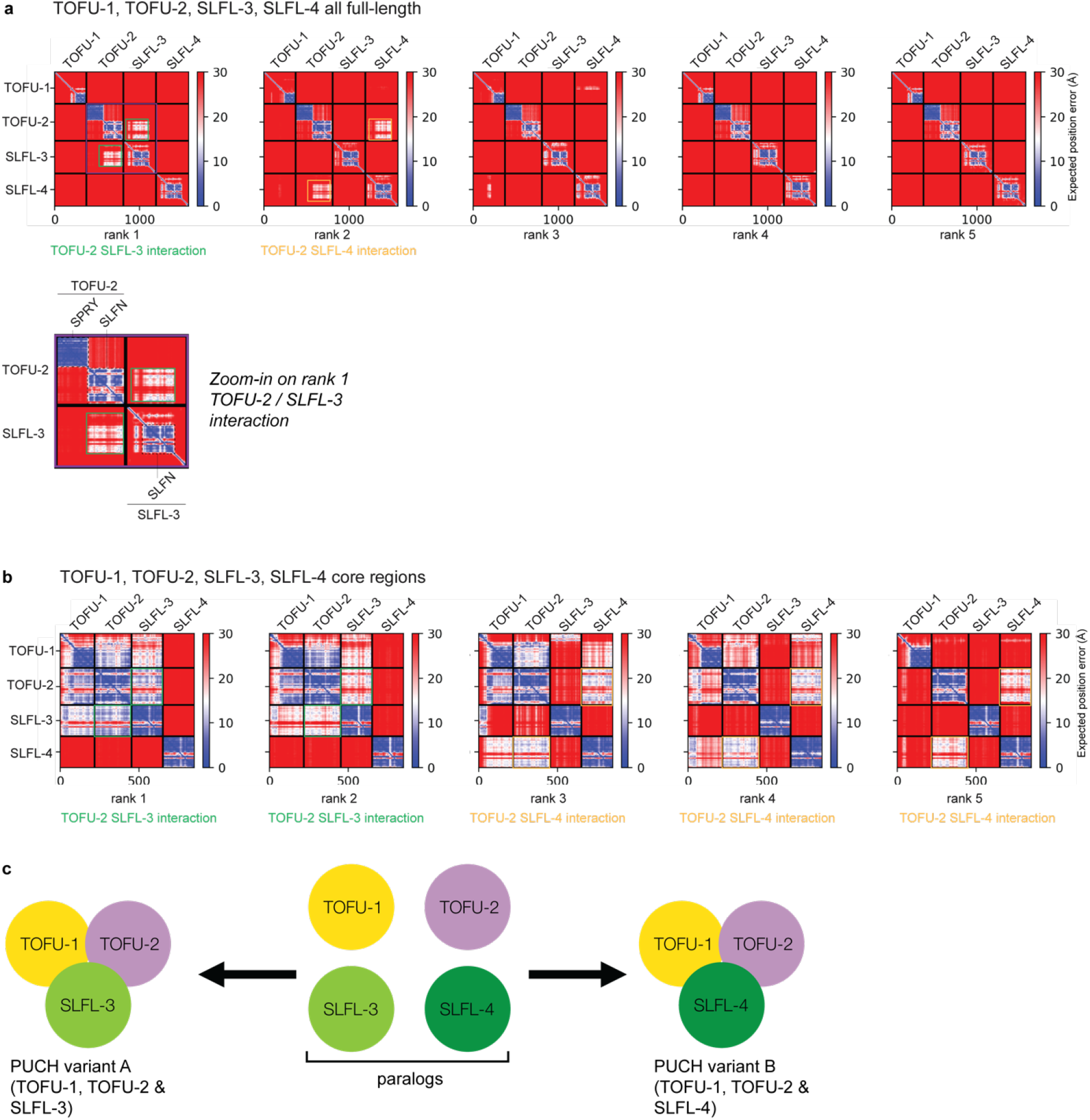
AlphaFold predicts a trimeric complex consisting of TOFU-1, TOFU-2 and either SLFL-3 or SLFL-4. **a**, Predicted alignment error (PAE) plots for the five models predicted by Alphafold for full-length TOFU-1, TOFU-2, SLFL-3 and SLFL-4. The zoom-in highlights the predicted interaction between the SLFN domains of TOFU-2 and SLFL-3, suggesting that the TOFU-2 SPRY domain is no involved in complex formation. The expected position error in angstroms (Å) is color coded where blue color indicates low PAE (high confidence) and red color indicates high PAE (low confidence). **b**, Predicted alignment error (PAE) plots for the five models predicted by Alphafold for core regions of TOFU-1, TOFU-1, SLFL-3 and SLFL-4. **c**, Schematic summary of the interaction results presented in panels **a** and **b**

**Fig. S4.**
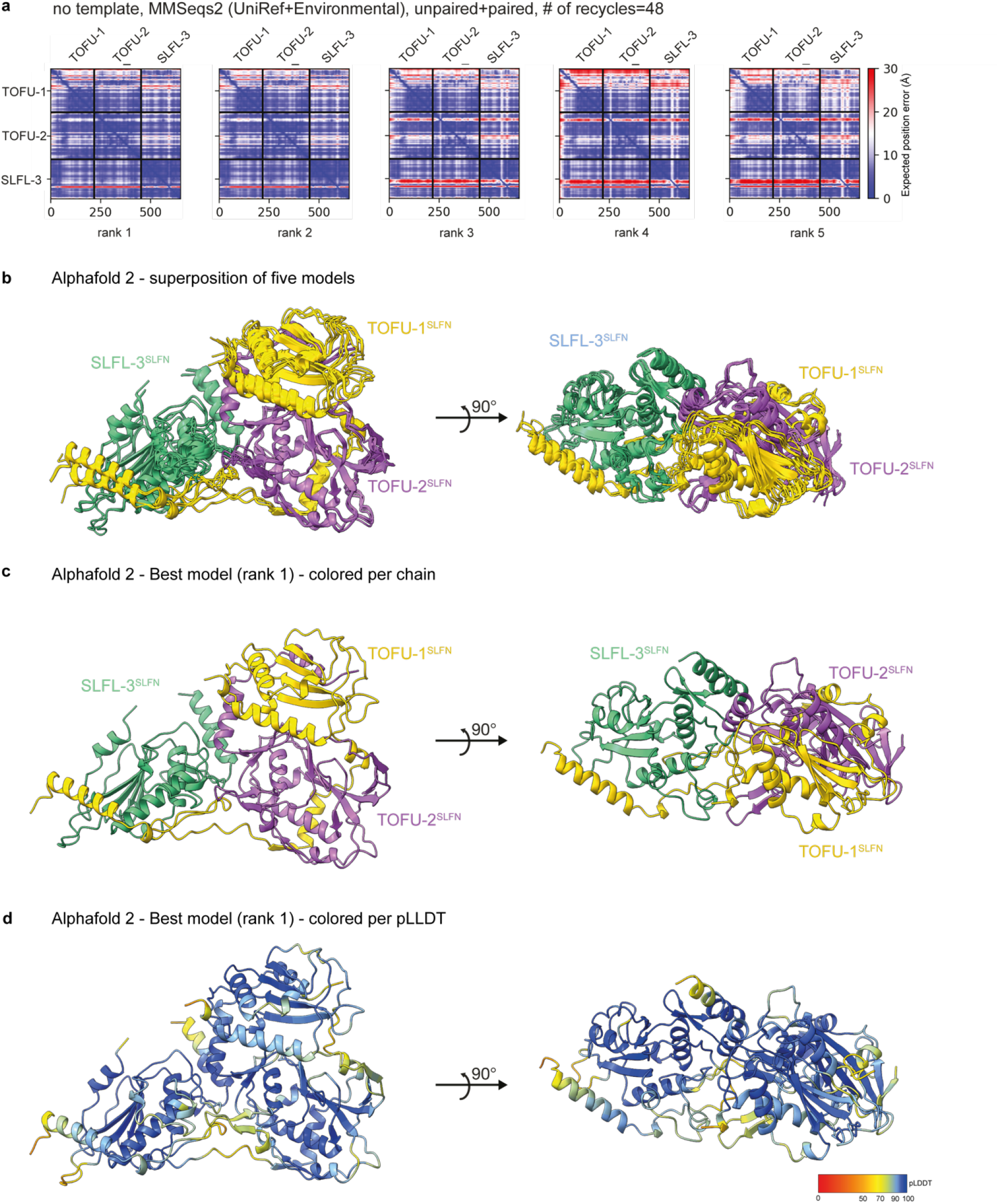
AlphaFold structure prediction of the trimeric complex consisting of TOFU-1, TOFU-2 and SLFL-3 shows convergence of models. **a**-**b**, AlphaFold predicts a trimeric complex consisting of TOFU-1, TOFU-2 and SLFL-3. TOFU-1 residues 156-373, TOFU-2 residues 200-433 and SLFL-3 residues 103-300 were used for the prediction. The predicted alignment error (PAE) plots are shown in (a), the five superposed models are shown as cartoon in (b). TOFU-1 is colored yellow, TOFU-2 purple and SLFL-3 green. The settings used for the prediction are shown on the top. The expected position error in angstroms (Å) is color coded where blue color indicates low PAE (high confidence) and red color indicates high PAE (low confidence). **c-d**, The best of the five predicted models is colored per chain (**c**) or per pLDDT score (**d**), which reports on the model confidence. Dark blue indicates very high, light blue confident, yellow low and orange very low model confidence.

**Fig. S5.**
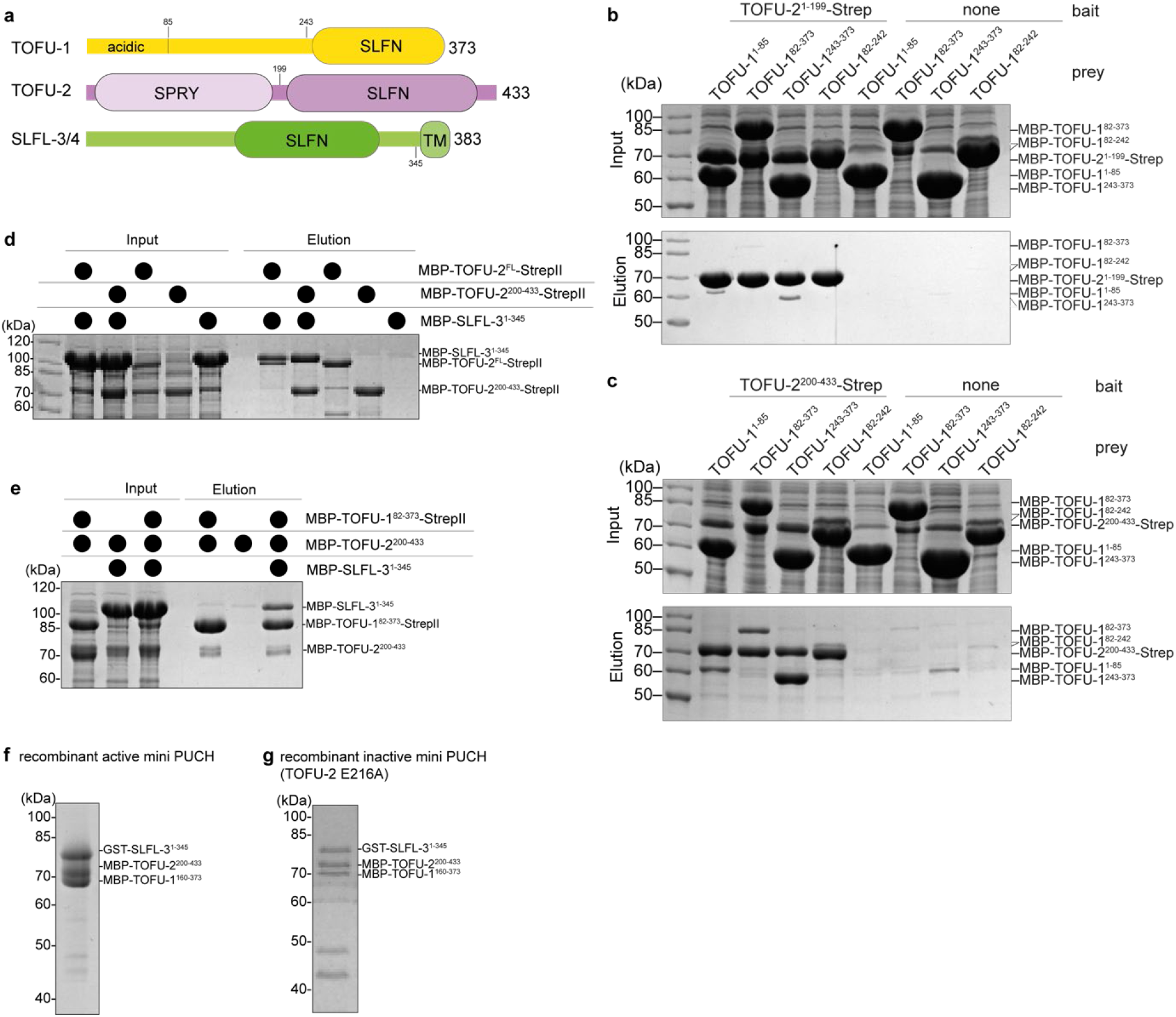
Verification of SLFL-interactions obtained by AlphaFold using recombinant proteins. **a**, Schematic domain organization of TOFU-1, TOFU-2 and SLFL-3. Lines indicate low-complexity regions and rounded rectangles indicate predicted folded domains. TM: transmembrane domain. **b-c**, A construct containing the TOFU-1 SLFN domain binds to the TOFU-2 SLFN domain while the TOFU-2 SPRY domain does not bind TOFU-1. Analysis of the interaction of different TOFU-1 constructs with the StrepII-tagged TOFU-2 SPRY domain in (b) and StrepII-tagged TOFU-2 SLFN domain in (c). The indicated constructs were co-expressed in *E. coli* and the StrepII-tagged bait was precipitated by Streptactin XT beads. Input and elution fractions were analyzed by SDS-PAGE followed by Coomassie staining. **d**, SLFL-3 interacts with the TOFU-2 SLFN domain. Analysis of the interaction of different StrepII-tagged TOFU-2 constructs with the SLFL-3. The indicated constructs were co-expressed in *E. coli* and the StrepII-tagged bait was precipitated by Streptactin XT beads. Input and elution fractions were analyzed by SDS-PAGE followed by Coomassie staining. **e**, TOFU-1, TOFU-2 and SLFL-3 form a trimeric complex. Different combinations of StrepII-tagged TOFU-1, TOFU-2 and SLFL-3 were co-expressed in *E. coli* and the StrepII-tagged bait was precipitated by Streptactin XT beads. Input and elution fractions were analyzed by SDS-PAGE followed by Coomassie staining. **f** and **g**, Recombinant, purified mini PUCH from *E. coli* in the active form (**f**) and inactive form TOFU-2 E216A (**g**).

**Fig. S6.**
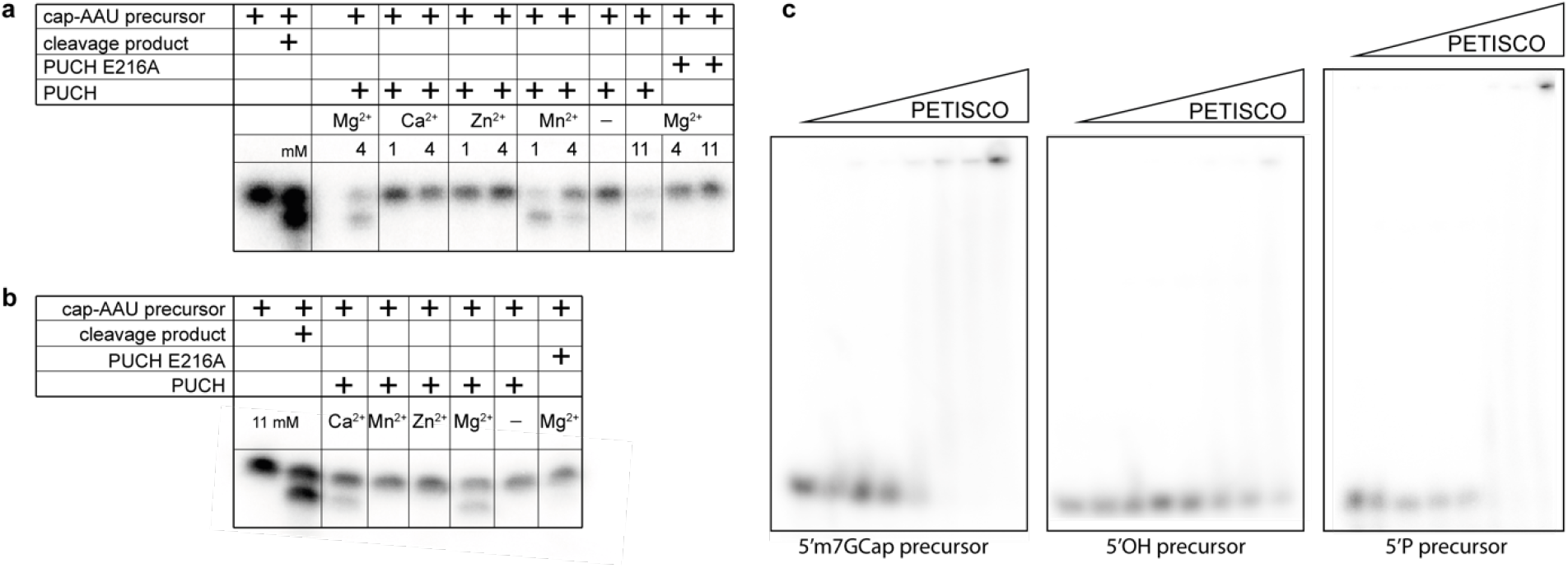
Divalent cation dependence of PUCH and RNA binding by PETISCO. **a-b**, Cleavage assays using GFP-IP material from BmN4 cell extracts and different divalent cations in various concentrations. **c**, Electrophoretic mobility shift assay between PETISCO complex and piRNA precursor with various 5’-ends.

**Fig. S7.**
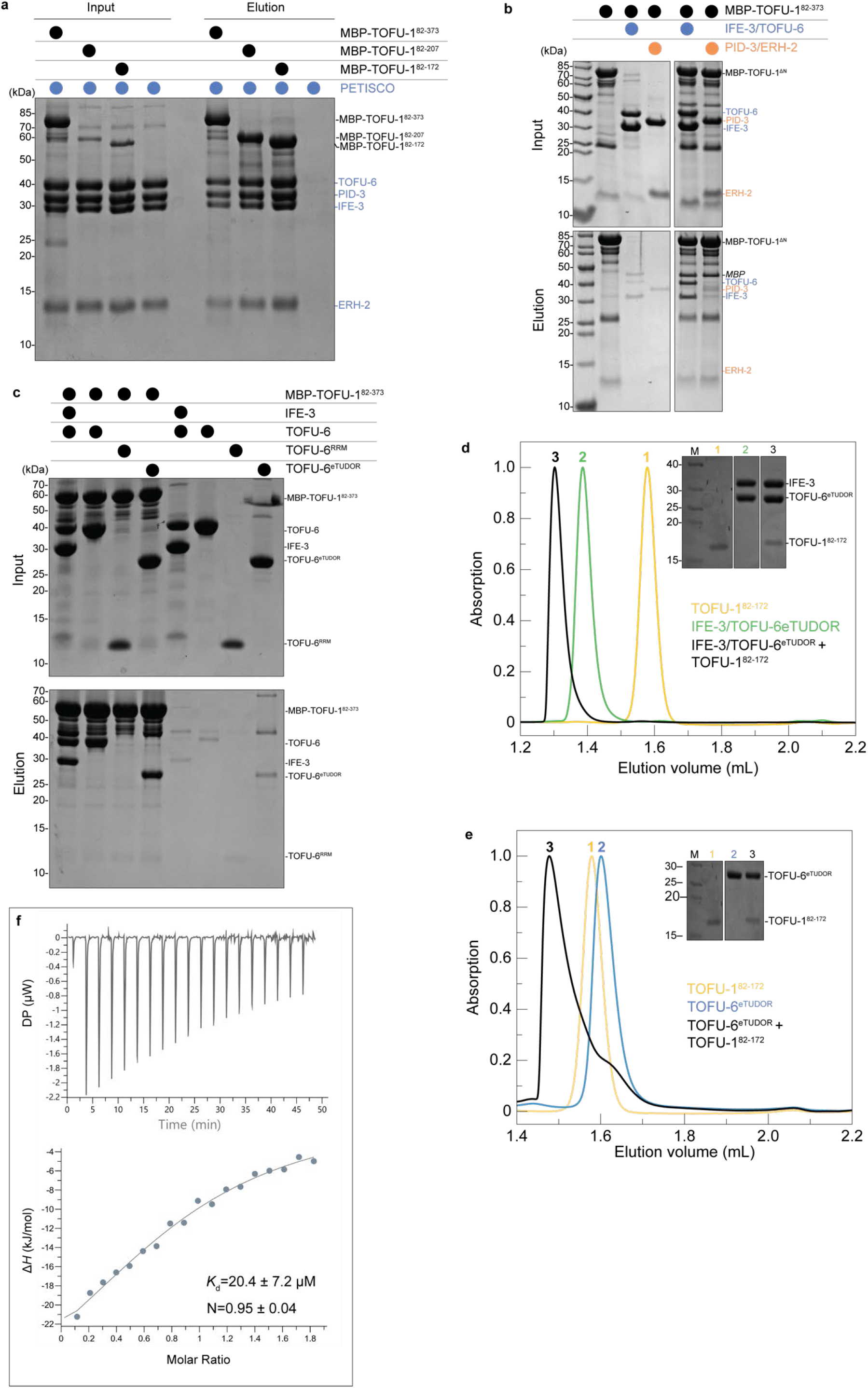
A peptide upstream of the TOFU-1 SLFN domain binds to the TOFU-6 eTUDOR domain. **a-c**, Analysis of the interaction of different TOFU-1 constructs with PETISCO and its subunits by amylose pull-down assays. Input and elution fractions were analyzed by SDS-PAGE followed by Coomassie staining. a, Various purified MBP-tagged TOFU-1 truncations were incubated with excess PETISCO and precipitated using amylose beads. b, Purified MBP-tagged TOFU-1^82-373^ was incubated with excess of the IFE-3/TOFU-6 and PID-3/ERH-2 subcomplexes precipitated using amylose beads. c, Purified MBP-tagged TOFU-1^82-373^ was incubated with excess of the IFE-3/TOFU-6 subcomplex, the TOFU-6 RRM and the TOFU-6 eTUDOR domain and precipitated using amylose beads. **d**, Purified IFE-3/TOFU-6^eTUDOR^ subcomplex, TOFU-1^72-182^ and a mixture thereof were subjected to size exclusion chromatography. Chromatograms: IFE-3/TOFU-6^eTUDOR^ (green), TOFU-1^72-182^ (yellow) and IFE-3/TOFU-6^eTUDOR^ + TOFU-1^72-182^ (black). The inset shows a Coomassie-stained SDS polyacrylamide gel of the peak fractions from size exclusion chromatography. **e**, Purified TOFU-6^eTUDOR^, TOFU-1^82-182^ and a mixture thereof were subjected to size exclusion chromatography. Chromatograms: TOFU-6^eTUDOR^ (blue), TOFU-1^182^ (yellow) and TOFU-6^eTUDOR^ + TOFU-1^182^ (black).The inset shows a Coomassie-stained SDS polyacrylamide gel of the peak fractions from size exclusion chromatography. Note: The chromatogram of TOFU-1^82-172^ (yellow) is shown for comparison and is the same as shown in panel **d**. Also, the lanes of the polyacrylamide gel are derived from the same gel in as in panel d; thus lane 1 and the marker are identical for panels **d** and **e.** **f**, Binding of TOFU-6^eTUDOR^ to TOFU-1^pep^ measured by ITC. The binding affinity (*K*_d_) and the stoichiometry (N) are the mean of three experiments and displayed error is the standard deviation. The experiment shows one of the three experiments as representative example.

**Fig. S8.**
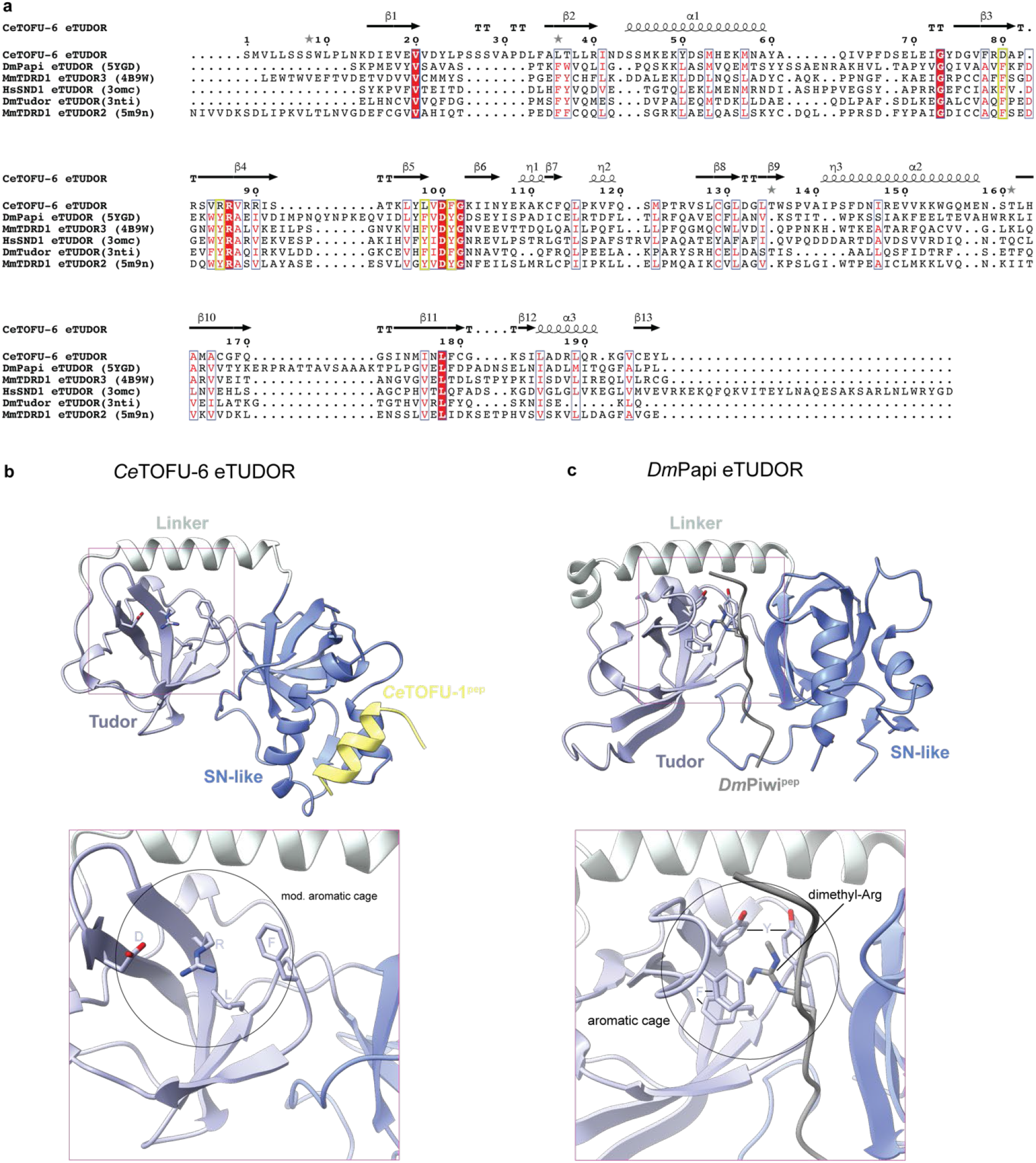
Comparison of the CeTOFU-6 eTUDOR domain to canonical eTUDOR domains. **a**, Structure-based sequence alignment of experimentally determined eTUDOR domains. Ce, *Caenorhabditis elegans*; Dm, *Drosophila melanogaster*; Mm, *Mus musculus* and Hs, *Homo sapiens*. The PDB ID is given in brackets. The secondary structure elements are indicated above the sequence and the four residues forming the aromatic cage in canonical eTUDOR domains are highlighted by yellow boxes. **b and c**, Crystal structures of the *C. elegans* TOFU-6^eTUDOR^–TOFU-1^pep^ and *D. melanogaster* PAPI^eTUDOR^–PIWI^pep^ complexes shown as cartoon. The eTUDOR domains are shown in different shades of blue, the TOFU-1^pep^ in yellow and the PIWI^pep^ containing the dimethyl-arginine residue in grey. The zoom-in view shows the region of the degenerated aromatic cage of TOFU-6^eTUDOR^ (**b**) and the canonical aromatic cage of PAPI^eTUDOR^ (**c**).

**Fig. S9.**
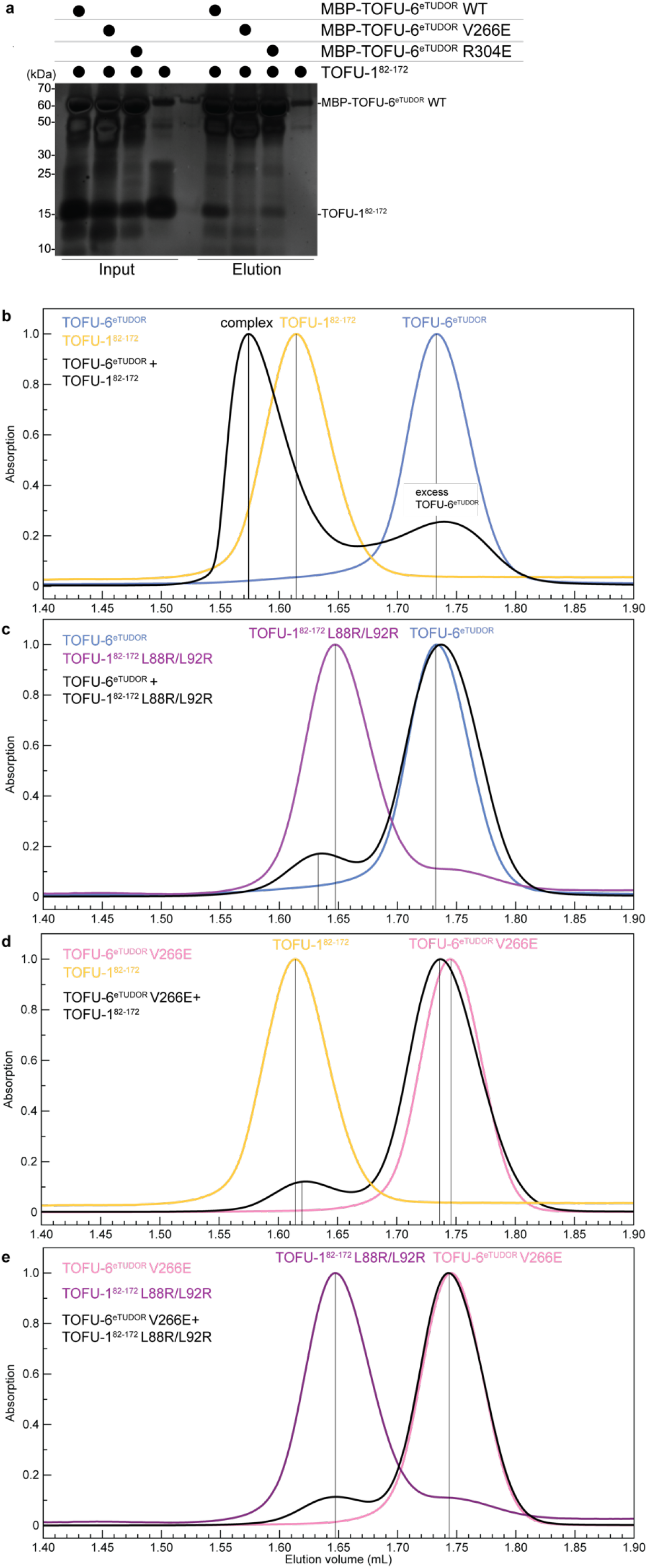
Structure-based analysis of the TOFU-6/TOFU-1 interaction. **a**, Pull-down assays with purified recombinant wild-type and mutant versions of MBP-TOFU-6^eTUDOR^ domain as bait and TOFU-1^82-172^ as prey. Input and elution fractions were analyzed by SDS-PAGE followed by silver staining. **b-e**, Analysis of the interaction between TOFU-1 and PETISCO by size exclusion chromatography. Purified recombinant wild-type and mutant versions of TOFU-6^eTUDOR^ and TOFU-1^82-172^ and mixtures thereof were subjected to size exclusion chromatography. Chromatograms: TOFU-6^eTUDOR^ (blue), TOFU-1^82-172^ (yellow), TOFU-6^eTUDOR^ V266E (pink), TOFU-1^82-172^ L88R/L92R (violet); the mixture of the respective proteins is always shown in black.

## References

1. Wells, J. N. & Feschotte, C. A Field Guide to Eukaryotic Transposable Elements. Annu Rev Genet 54, 539–561 (2020).

2. Ozata, D. M., Gainetdinov, I., Zoch, A., O’Carroll, D. & Zamore, p. D. PIWI-interacting RNAs: small RNAs with big functions. Nat Rev Genet 20, 89–108 (2019).

3. Czech, B. et al. piRNA-Guided Genome Defense: From Biogenesis to Silencing. Annu Rev Genet 52, 131–157 (2018).

4. Cordeiro Rodrigues, R. J. et al. PETISCO is a novel protein complex required for 21U RNA biogenesis and embryonic viability. Genes Dev 33, (2019).

5. Jo, U. & Pommier, Y. Structural, molecular, and functional insights into Schlafen proteins. Exp Mol Med 54, 730–738 (2022).

6. Cosby, R. L., Chang, N.-C. & Feschotte, C. Host-transposon interactions: conflict, cooperation, and cooption. Genes Dev 33, 1098–1116 (2019).

7. Wu, P.-H. & Zamore, p. D. Defining the functions of PIWI-interacting RNAs. Nat Rev Mol Cell Biol 22, 239–240 (2021).

8. Luteijn, M. J. & Ketting, R. F. PIWI-interacting RNAs: From generation to transgenerational epigenetics. Nat Rev Genet 14, (2013).

9. Voigt, F. et al. Crystal structure of the primary piRNA biogenesis factor Zucchini reveals similarity to the bacterial PLD endonuclease Nuc. RNA 18, 2128–34 (2012).

10. Ipsaro, J. J., Haase, A. D., Knott, S. R., Joshua-Tor, L. & Hannon, G. J. The structural biochemistry of Zucchini implicates it as a nuclease in piRNA biogenesis. Nature 491, 279–83 (2012).

11. Izumi, N., Shoji, K., Suzuki, Y., Katsuma, S. & Tomari, Y. Zucchini consensus motifs determine the mechanism of pre-piRNA production. Nature 578, 311–316 (2020).

12. Nishimasu, H. et al. Structure and function of Zucchini endoribonuclease in piRNA biogenesis. Nature 491, 284–7 (2012).

13. Mohn, F., Handler, D. & Brennecke, J. Noncoding RNA. piRNA-guided slicing specifies transcripts for Zucchini-dependent, phased piRNA biogenesis. Science 348, 812–817 (2015).

14. Han, B. W., Wang, W., Li, C., Weng, Z. & Zamore, p. D. Noncoding RNA. piRNA-guided transposon cleavage initiates Zucchini-dependent, phased piRNA production. Science 348, 817–21 (2015).

15. Tang, W., Tu, S., Lee, H.-C., Weng, Z. & Mello, C. C. The RNase PARN-1 Trims piRNA 3’ Ends to Promote Transcriptome Surveillance in C. elegans. Cell 164, 974–84 (2016).

16. Montgomery, T. A. et al. PIWI associated siRNAs and piRNAs specifically require the Caenorhabditis elegans HEN1 ortholog henn-1. PLoS Genet 8, (2012).

17. Billi, A. C. et al. The caenorhabditis elegans HEN1 Ortholog, HENN-1, methylates and stabilizes select subclasses of germline small RNAs. PLoS Genet 8, (2012).

18. Kamminga, L. M. et al. Differential impact of the HEN1 homolog HENN-1 on 21U and 26G RNAs in the germline of Caenorhabditis elegans. PLoS Genet 8, (2012).

19. Wang, G. & Reinke, V. A C. elegans Piwi, PRG-1, regulates 21U-RNAs during spermatogenesis. Curr Biol 18, 861–7 (2008).

20. Batista, P. J. et al. PRG-1 and 21U-RNAs interact to form the piRNA complex required for fertility in C. elegans. Mol Cell 31, 67–78 (2008).

21. Das, P. P. et al. Piwi and piRNAs Act Upstream of an Endogenous siRNA Pathway to Suppress Tc3 Transposon Mobility in the Caenorhabditis elegans Germline. Mol Cell 31, (2008).

22. Bagijn, M. P. et al. Function, targets, and evolution of Caenorhabditis elegans piRNAs. Science 337, 574–578 (2012).

23. Ruby, J. G. et al. Large-scale sequencing reveals 21U-RNAs and additional microRNAs and endogenous siRNAs in C. elegans. Cell 127, 1193–207 (2006).

24. Gu, W. et al. CapSeq and CIP-TAP identify Pol II start sites and reveal capped small RNAs as C. elegans piRNA precursors. Cell 151, 1488–500 (2012).

25. Blumenthal, T. Trans-splicing and operons in C. elegans. WormBook 1–11 (2012) doi:10.1895/wormbook.1.5.2.

26. Weick, E.-M. et al. PRDE-1 is a nuclear factor essential for the biogenesis of Ruby motif-dependent piRNAs in C. elegans. Genes Dev 28, 783–96 (2014).

27. Weng, C. et al. The USTC co-opts an ancient machinery to drive piRNA transcription in C. elegans. Genes Dev 33, 90–102 (2019).

28. Perez-Borrajero, C. et al. Structural basis of PETISCO complex assembly during piRNA biogenesis in C. elegans. Genes Dev 35, 1304–1323 (2021).

29. Wang, X. et al. Molecular basis for PICS-mediated piRNA biogenesis and cell division. Nat Commun 12, (2021).

30. Zeng, C. et al. Functional Proteomics Identifies a PICS Complex Required for piRNA Maturation and Chromosome Segregation. Cell Rep 27, 3561-3572.e3 (2019).

31. Goh, W.-S. S. et al. A genome-wide RNAi screen identifies factors required for distinct stages of C. elegans piRNA biogenesis. Genes Dev 28, 797–807 (2014).

32. Zimmermann, L. et al. A Completely Reimplemented MPI Bioinformatics Toolkit with a New HHpred Server at its Core. J Mol Biol 430, 2237–2243 (2018).

33. Jumper, J. et al. Highly accurate protein structure prediction with AlphaFold. Nature 596, 583–589 (2021).

34. Yang, J.-Y. et al. Structure of Schlafen13 reveals a new class of tRNA/rRNA-targeting RNase engaged in translational control. Nat Commun 9, 1165 (2018).

35. Metzner, F. J., Huber, E., Hopfner, K.-P. & Lammens, K. Structural and biochemical characterization of human Schlafen 5. Nucleic Acids Res 50, 1147–1161 (2022).

36. Garvie, C. W. et al. Structure of PDE3A-SLFN12 complex reveals requirements for activation of SLFN12 RNase. Nat Commun 12, (2021).

37. Varadi, M. et al. AlphaFold Protein Structure Database: massively expanding the structural coverage of protein-sequence space with high-accuracy models. Nucleic Acids Res 50, D439–D444 (2022).

38. Watanabe, T. et al. MITOPLD is a mitochondrial protein essential for nuage formation and piRNA biogenesis in the mouse germline. Dev Cell 20, 364–75 (2011).

39. Huang, H. et al. piRNA-associated germline nuage formation and spermatogenesis require MitoPLD profusogenic mitochondrial-surface lipid signaling. Dev Cell 20, 376–87 (2011).

40. Evans, R. et al. Protein complex prediction with AlphaFold-Multimer. bioRxiv 2021.10.04.463034 (2022) doi:10.1101/2021.10.04.463034.

41. Metzner, F. J. et al. Mechanistic understanding of human SLFN11. Nat Commun 13, 5464 (2022).

42. Yang, W. Nucleases: diversity of structure, function and mechanism. Q Rev Biophys 44, 1–93 (2011).

43. de Albuquerque, B. F. M. et al. PID-1 is a novel factor that operates during 21U-RNA biogenesis in Caenorhabditis elegans. Genes Dev 28, (2014).

44. Gan, B., Chen, S., Liu, H., Min, J. & Liu, K. Structure and function of eTudor domain containing TDRD proteins. Crit Rev Biochem Mol Biol 54, 119–132 (2019).

45. Chen, C., Nott, T. J., Jin, J. & Pawson, T. Deciphering arginine methylation: Tudor tells the tale. Nat Rev Mol Cell Biol 12, 629–642 (2011).

46. Bell, T. A. et al. The paternal gene of the DDK syndrome maps to the Schlafen gene cluster on mouse chromosome 11. Genetics 172, 411–23 (2006).

47. Bartonicek, N. et al. The retroelement Lx9 puts a brake on the immune response to virus infection. Nature 608, 757–765 (2022).

48. Li, M. et al. Codon-usage-based inhibition of HIV protein synthesis by human schlafen 11. Nature 491, 125–8 (2012).

49. Eaglesham, J. B., McCarty, K. L. & Kranzusch, p. J. Structures of diverse poxin cGAMP nucleases reveal a widespread role for cGAS-STING evasion in host-pathogen conflict. Elife 9, (2020).

50. Gubser, C. et al. Camelpox virus encodes a schlafen-like protein that affects orthopoxvirus virulence. J Gen Virol 88, 1667–1676 (2007).

51. Glover, M. L. et al. NONU-1 Encodes a Conserved Endonuclease Required for mRNA Translation Surveillance. Cell Rep 30, 4321-4331.e4 (2020).

52. Brenner, S. The genetics of Caenorhabditis elegans. Genetics 77, 71–94 (1974).

53. Schweinsberg, P. J. & Grant, B. D. C. elegans gene transformation by microparticle bombardment. WormBook 1–10 (2013) doi:10.1895/WORMBOOK.1.166.1.

54. Arribere, J. A. et al. Efficient marker-free recovery of custom genetic modifications with CRISPR/Cas9 in Caenorhabditis elegans. Genetics 198, 837–846 (2014).

55. Rappsilber, J., Mann, M. & Ishihama, Y. Protocol for micro-purification, enrichment, pre-fractionation and storage of peptides for proteomics using StageTips. Nature Protocols 2007 2:8 2, 1896–1906 (2007).

56. Cox, J. & Mann, M. Quantitative, High-Resolution Proteomics for Data-Driven Systems Biology. https://doi.org/10.1146/annurev-biochem-061308-093216 80, p273–299 (2011).

57. Cox, J. & Mann, M. MaxQuant enables high peptide identification rates, individualized p.p.b.-range mass accuracies and proteome-wide protein quantification. Nature Biotechnology 2008 26:12 26, 1367–1372 (2008).

58. Gorrec, F. The MORPHEUS protein crystallization screen. J Appl Crystallogr 42, 1035–1042 (2009).

59. Newman, J. et al. Towards rationalization of crystallization screening for small-to medium-sized academic laboratories: the PACT/JCSG+ strategy. Acta Crystallogr D Biol Crystallogr 61, 1426–31 (2005).

60. Vonrhein, C. et al. Data processing and analysis with the autoPROC toolbox. Acta Crystallogr D Biol Crystallogr 67, 293–302 (2011).

61. Kabsch, W. XDS. Acta Crystallogr D Biol Crystallogr 66, 125–32 (2010).

62. Evans, P. R. & Murshudov, G. N. How good are my data and what is the resolution? Acta Crystallogr D Biol Crystallogr 69, 1204–14 (2013).

63. McCoy, A. J. et al. Phaser crystallographic software. J Appl Crystallogr 40, 658–674 (2007).

64. Liebschner, D. et al. Macromolecular structure determination using X-rays, neutrons and electrons: recent developments in Phenix. Acta Crystallogr D Struct Biol 75, 861–877 (2019).

65. Cowtan, K. The Buccaneer software for automated model building. 1. Tracing protein chains. Acta Crystallogr D Biol Crystallogr 62, 1002–11 (2006).

66. Emsley, P., Lohkamp, B., Scott, W. G. & Cowtan, K. Features and development of Coot. Acta Crystallogr D Biol Crystallogr 66, 486–501 (2010).

67. Afonine, P. v et al. Towards automated crystallographic structure refinement with phenix.refine. Acta Crystallogr D Biol Crystallogr 68, 352–67 (2012).

68. Williams, C. J. et al. MolProbity: More and better reference data for improved all-atom structure validation. Protein Sci 27, 293–315 (2018).

69. Joosten, R. P., Joosten, K., Murshudov, G. N. & Perrakis, A. PDB_REDO: constructive validation, more than just looking for errors. Acta Crystallogr D Biol Crystallogr 68, 484–96 (2012).

70. Pettersen, E. F. et al. UCSF ChimeraX: Structure visualization for researchers, educators, and developers. Protein Sci 30, 70–82 (2021).

71. Mirdita, M. et al. ColabFold: making protein folding accessible to all. Nat Methods 19, 679–682 (2022).

72. Perez-Riverol, Y. et al. The PRIDE database resources in 2022: a hub for mass spectrometry-based proteomics evidences. Nucleic Acids Res 50, D543–D552 (2022).

