## Supplementary material for "piRNA processing by a trimeric Schlafen-domain nuclease": structure data

**Supplemental Table S3**
**Data collection and refinement statistics (molecular replacement)**

| | TOFU-6$^{eTUDOR}$/TOFU-1$^{pep}$ complex (PDB code: 8BY5) |
|---|---|
| **Data collection** | 26th September 2021 |
| Space group | C 1 2 1 |
| Cell dimensions | |
| *a*, *b*, *c* (Å) | 59.12 68.13 55.17 |
| α, β, γ (°) | 90.00, 103.03, 90.00 |
| Resolution (Å) | 34.06 – 1.73 (1.79 – 1.73)† |
| $R_{merge}$ | 0.056 (0.479) |
| $I / \sigma I$ | 17.8 (2.4) |
| Completeness (%) | 93.5 (66.7) |
| Redundancy | 5.6 (3.9) |
| CC (1/2) | 99.9 (79.4) |
| **Refinement** | |
| Resolution (Å) | 1.73 |
| No. unique reflections | 20818 (1469) |
| $R_{work}$ | 0.162 (0.264) |
| $R_{free}$ | 0.207 (0.313) |
| No. atoms | |
| Protein | 1660 |
| Ligand/ion | 18 |
| Water | 205 |
| *B*-factors | |
| Protein | 30.3 |
| Ligand/ion | 43.5 |
| Water | 34.3 |
| R.m.s. deviations | |
| Bond lengths (Å) | 0.016 |
| Bond angles (°) | 1.18 |

*Number of xtals for each structure:1 should be noted in footnote.
†Values in parentheses are for highest-resolution shell.
